# The conserved elongation factor Spn1 is required for normal transcription, histone modifications, and splicing in *Saccharomyces cerevisiae*

**DOI:** 10.1101/2020.07.01.182956

**Authors:** Natalia I. Reim, James Chuang, Dhawal Jain, Burak H. Alver, Peter J. Park, Fred Winston

## Abstract

Spn1/Iws1 is a conserved protein involved in transcription and chromatin dynamics, yet its general *in vivo* requirement for these functions is unknown. Using a Spn1 depletion system in *S. cerevisiae,* we demonstrate that Spn1 broadly influences several aspects of gene expression on a genome-wide scale. We show that Spn1 is globally required for normal mRNA levels and for normal splicing of ribosomal protein transcripts. Furthermore, Spn1 maintains the localization of H3K36 and H3K4 methylation across the genome and is required for normal histone levels at highly expressed genes. Finally, we show that the association of Spn1 with the transcription machinery is strongly dependent on its binding partner, Spt6, while the association of Spt6 and Set2 with transcribed regions is partially dependent on Spn1. Taken together, our results show that Spn1 affects multiple aspects of gene expression and provide additional evidence that it functions as a histone chaperone *in vivo*.

## Introduction

Spn1/Iws1 (hereafter called Spn1) is an evolutionarily conserved protein that is essential for viability in *Saccharomyces cerevisiae* and human cells and plays a role in mammalian and plant development^1–6^. Spn1 was first identified in *S. cerevisiae* as a protein that interacts physically and genetically with factors involved in transcription^1,7–9^, and it was later found to be part of the general RNA polymerase II (RNAPII) elongation complex on actively transcribed genes^10^. Several studies have suggested that Spn1 is involved in transcription and co-transcriptional processes, including transcriptional activation^1,11,12^, elongation^1,8–10^, pre-mRNA processing^3,5,13–15^, and mRNA nuclear export^3,13,14^. The recent demonstration that Spn1 binds histones and assembles nucleosomes *in vitro* suggested that Spn1 functions as a histone chaperone^16^.

Spn1 directly interacts with Spt6^17,18^, another essential histone chaperone that associates with elongating RNAPII^10,19,20^, and some evidence suggests that the two factors contribute to a common function. For example, some *spn1* and *spt6* mutations exhibit similar genetic interactions with mutations in genes encoding other factors involved in transcription and chromatin dynamics^21,22^. In addition, combining *spn1* and *spt6* mutations or disrupting the Spn1-Spt6 interaction by mutation results in synthetic lethality, poor growth, or phenotypes associated with defective transcription and chromatin organization (^17,18,23^ and our unpublished results). However, there is also evidence for separate functions of Spn1 and Spt6 based on distinct recruitment patterns to transcribed chromatin^10^ and different functional requirements at the yeast *CYC1* gene^1^.

Studies have suggested a role for Spn1 in histone H3K36 methylation. In mammalian cells, it was proposed that Spn1 is required for H3K36 trimethylation by mediating the interaction between Spt6 and the histone methyltransferase SETD2 during elongation by RNAPII^5,14^. In yeast, while Spt6 is clearly required for Set2-dependent H3K36 methylation^24–30^, the requirement for Spn1 is less certain, as mutations disrupting the Spn1-Spt6 interaction did not cause a complete loss of H3K36me3 and H3K36me2 (^30^ and our unpublished results).

A functional connection between Spn1 and H3K36 methylation in yeast came from recent genetic evidence that showed that null mutations in genes encoding Set2 or components of the Rpd3S histone deacetylase complex (*set2Δ*, *rpd3Δ*, *eaf3Δ*, *rco1Δ*) suppress the temperature sensitive phenotype of a *spn1* mutation, *spn1-K192N*^22^. Rpd3S is recruited and activated by H3K36me2/3 and the Ser5-phosphorylated RNAPII CTD^24,31–35^. This suppression of *spn1-K192N*^22^ suggested that one role of Spn1 is to oppose the functions of the Rpd3S histone deacetylase complex and, furthermore, that increased histone acetylation can compensate for impairment of Spn1 function.

In contrast to the activation of Rpd3S by H3K36 methylation, Set1-dependent H3K4 methylation has been proposed to oppose the function of Rpd3S, similar to the role proposed for Spn1^22,36^. By this model, H3K4me3 would inhibit deacetylation by Rpd3S in the region immediately downstream of promoters, where H3K4me3 is primarily found^37–41^. In line with the idea that H3K36 methylation and H3K4 methylation have opposing functions, studies showed that loss of H3K4 methylation exacerbated the Ts^−^ phenotype of *spn1-K192N,* while increased levels of H3K4 methylation suppressed *spn1-K192*N^21,22^. In summary, genetic analysis suggests that the Set2-Rpd3S pathway opposes the function of Spn1, while Set1-dependent H3K4 methylation promotes it.

In this work, we have performed experiments to understand the requirement for Spn1 in transcription and co-transcriptional processes *in vivo.* Using an inducible Spn1 depletion system and RNA-seq, we found that loss of Spn1 results in widespread changes in mRNA levels. Furthermore, using ChIP-seq, we found that Spn1 depletion causes redistribution of H3K36me3, H3K36me2, and H3K4me3 genome-wide, as well as histone loss at highly transcribed genes. Additional experiments showed that Spt6 is crucial for the association of Spn1 with RNAPII, while Spn1 is not strongly required for the interaction of Spt6 with RNAPII but rather plays a role in optimal Spt6 recruitment to chromatin. Finally, our results showed that Spn1 is required for efficient splicing of ribosomal protein transcripts. Together, our results reveal that Spn1 has widespread roles in the control of transcription and chromatin.

## Results

### An auxin-inducible degron system efficiently depletes Spn1 in *S. cerevisiae*

To study the requirement for the essential protein Spn1, we constructed an *S. cerevisiae* strain for the depletion of Spn1 using an auxin-inducible degron (see Methods). Treatment of this strain with the auxin 3-indoleacetic acid (IAA) for 90 minutes (Spn1-depleted condition) resulted in the reduction of Spn1 protein levels to about 8% of those observed when the strain was treated with DMSO (non-depleted condition) (Fig. 1a). Importantly, these conditions did not impair viability (Supplementary Fig. 1a).

**Figure 1.**
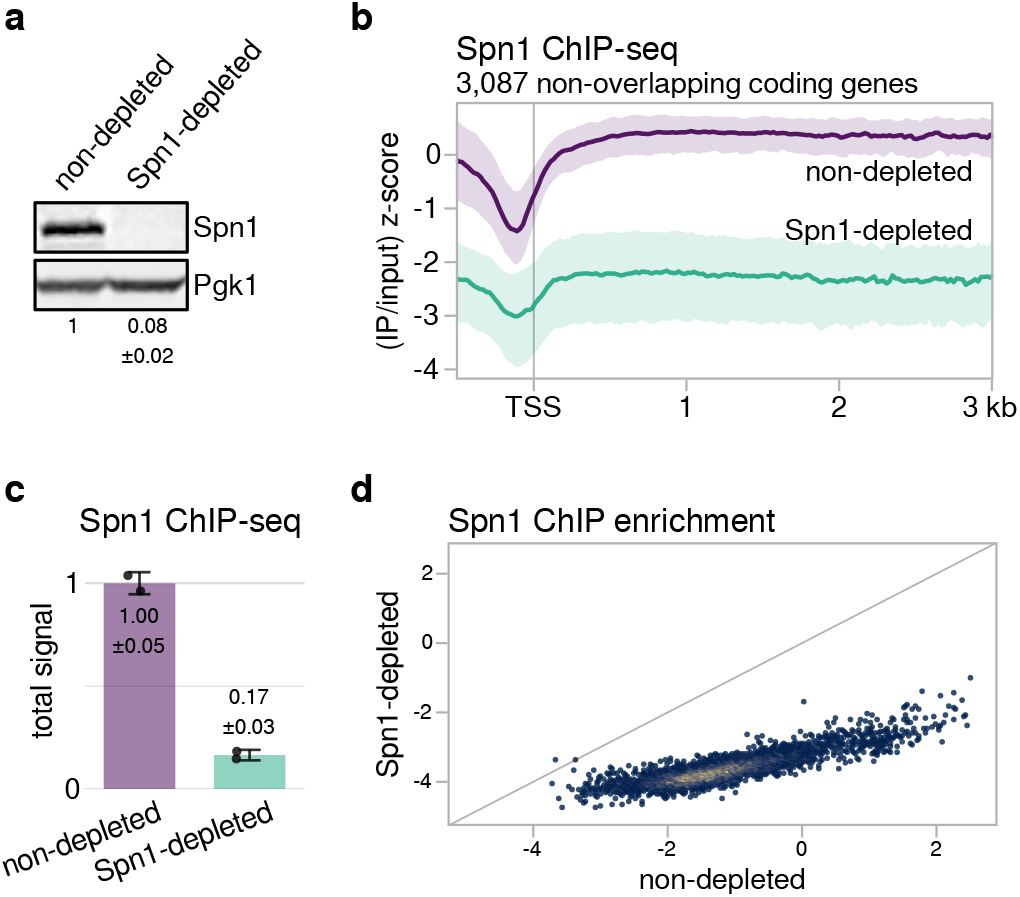
An auxin-inducible degron efficiently depletes Spn1 from chromatin. **a** Western blot showing levels of Spn1 protein in the Spn1 degron strain after 90 minutes of treatment with DMSO (non-depleted) or 25 μM IAA (Spn1-depleted). Pgk1 was used as a loading control. The values below the blots indicate the mean ± standard deviation of normalized Spn1 abundance quantified from two biological replicates. **b** Average Spn1 ChIP enrichment over 3,087 non-overlapping, verified coding genes aligned by TSS in non-depleted and Spn1-depleted conditions. For each gene, the spike-in normalized ratio of IP over input signal is standardized to the mean and standard deviation of the non-depleted signal over the region, resulting in standard scores that allow genes with varied expression levels to be compared on the same scale. The solid line and shading are the median and interquartile range of the mean standard score over two replicates. **c** Barplot showing total levels of Spn1 on chromatin in non-depleted and Spn1-depleted conditions, estimated from ChIP-seq spike-in normalization factors (see Methods). The values and error bars for each condition indicate the mean ± standard deviation of two replicates. **d** Scatterplot showing Spn1 ChIP enrichment in Spn1-depleted versus non-depleted conditions for 5,091 verified coding genes. Enrichment values are the relative log_2_ enrichment of IP over input.

To verify that Spn1 depletion reduced the amount of Spn1 on chromatin, we performed chromatin immunoprecipitation and high-throughput sequencing (ChIP-seq) for Spn1 in Spn1-depleted and non-depleted cells, using *S. pombe* chromatin as a spike-in control to allow the detection of global changes (see Methods). In agreement with a previous study^10^, Spn1 in non-depleted cells was primarily enriched over gene bodies (Fig. 1b) at levels strongly correlated with levels of the RNAPII subunit Rpb1 (Supplementary Fig. 1b). Additionally, we noted that the ratio of Spn1 to Rpb1 over coding genes decreased slightly with increasing Rpb1 levels (Supplementary Fig. 1c). In Spn1-depleted cells, the total Spn1 ChIP signal was approximately 17% that of non-depleted cells (Fig. 1c). The reduction in Spn1 levels was observed for virtually all coding genes (Fig. 1d) and occurred uniformly over gene bodies (Fig. 1b). Taken together, our results show that the auxin-inducible degron system effectively depletes Spn1 from chromatin genome-wide and allows for interrogation of its essential functions during transcription.

### Depletion of Spn1 causes a widespread reduction of mRNA levels

Although there is evidence that Spn1 is involved in transcription, the genome-wide requirement for Spn1 in regulating mRNA levels was unknown. To investigate this, we performed RNA sequencing (RNA-seq) of Spn1-depleted and non-depleted cells, using *S. pombe* cells as a spike-in control. The major effect that we observed was a widespread decrease in mRNA abundance in Spn1-depleted versus non-depleted cells, with a median change to around 73% of non-depleted transcript levels and over 1500 mRNAs detected as differentially reduced (Fig. 2a). We also detected 33 mRNAs with increased levels (Fig. 2a). By analyzing RNA-seq data from additional control conditions, we verified that the changes in transcript abundance were caused primarily by Spn1 depletion rather than by the presence of the degron tag on Spn1, the expression of degron system components, or the effects of auxin on yeast cells (see Methods; Supplementary Fig. 2a). We verified the differential expression of seven genes by RT-qPCR (Supplementary Fig. 2b), including the strong induction of *SER3* previously observed in *spn1* mutants^42^.

**Figure 2.**
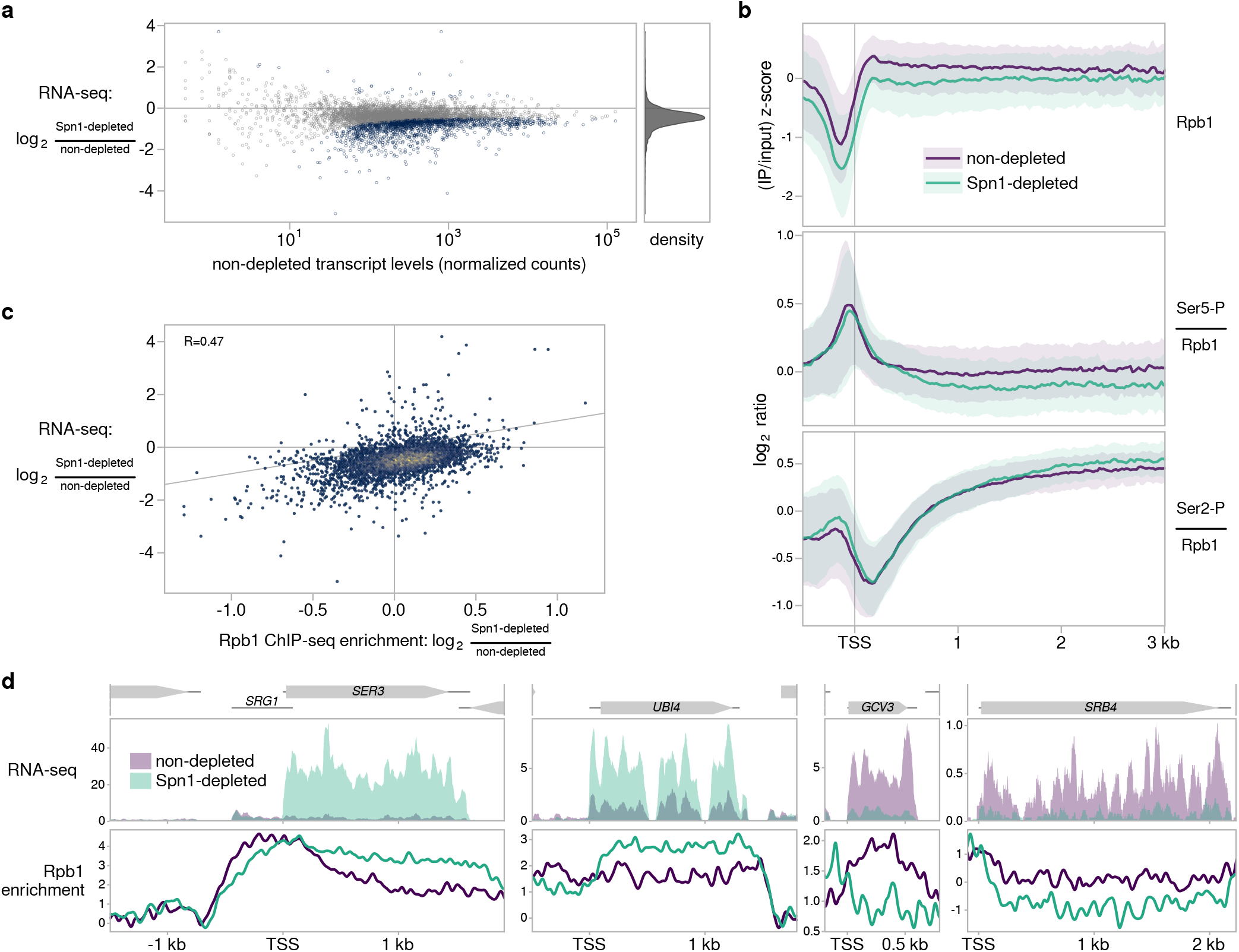
Effects of Spn1 depletion on mRNA levels and RNAPII. **a** Scatterplot showing changes in transcript abundance as measured by RNA-seq upon Spn1 depletion versus non-depleted transcript abundance for 5,091 verified coding transcripts. Transcripts with significant changes are colored blue, and the relative density of fold-change values is shown in the right panel. **b** Top panel: Average Rpb1 ChIP enrichment over 3,087 non-overlapping, verified coding genes in non-depleted and Spn1-depleted conditions. Spike-in normalized standard scores are calculated as in Fig. 1b. The solid line and shading are the median and interquartile range of the mean standard score over the two replicates generated from the same chromatin preparations used to perform ChIP-seq for Rpb1-Ser5-P and Rpb1-Ser2-P. Bottom two panels: Average Rpb1-normalized Rpb1-Ser5-P and Rpb1-Ser2-P ChIP enrichment in non-depleted and Spn1-depleted conditions for the same genes. The solid line and shading are the median and interquartile range of the mean spike-in normalized ratio over two replicates. **c** Scatterplot showing change in transcript abundance upon Spn1 depletion versus change in Rpb1 enrichment for 5,091 verified coding genes. The line y=x is drawn for comparison, and the Pearson correlation coefficient is shown. **d** Spike-in normalized sense strand RNA-seq coverage and spike-in normalized Rpb1 ChIP enrichment over four genes, two with elevated mRNA levels and two with decreased mRNA levels upon Spn1 depletion.

In response to various environmental stresses, yeast cells are known to alter the transcript levels of approximately 870 environmental stress response (ESR) genes^43^. To look for a possible relationship between Spn1 depletion and the ESR, we performed gene set enrichment analysis. Our results revealed an association between Spn1-dependent transcriptional changes and the ESR (Supplementary Fig. 2c), consistent with the fact that Spn1 is essential for viability. However, the transcriptional changes in response to Spn1 depletion are more global in scope than the ESR, do not correlate with the gene expression changes of any individual ESR stress (data not shown), and do not correlate with a related transcriptional signature of slow growth^44^ (Supplementary Fig. 2d). This suggests that the Spn1-dependent changes we observe are not only explained by stress responses.

Impaired function of the Spn1 binding partner Spt6 causes widespread derepression of antisense transcripts^19,26,45–48^. Therefore, we were interested to see whether a similar effect was observed after Spn1 depletion. In contrast to *spt6* mutants, we detected only four antisense transcripts with significantly increased levels in Spn1-depleted cells (Supplementary Fig. 2e). Though this may be a slight underestimate due to the difficulty of annotating lowly expressed transcripts, it is clear that antisense transcripts are much more limited for Spn1 depletion as compared to *spt6* mutants.

To test whether the widespread changes in sense transcript levels observed after Spn1 depletion might be driven by changes in the level or state of RNAPII, we performed ChIP-seq for the largest RNAPII subunit, Rpb1, as well for two major Rpb1 C-terminal domain (CTD) post-translational modifications: serine 5 phosphorylation (Ser5-P) and serine 2 phosphorylation (Ser2-P). Our results show that the relative distribution of Rpb1 over transcribed regions was not greatly affected by Spn1 depletion (Fig. 2b). However, we observed that the modest changes in Rpb1 levels in Spn1-depleted cells correlate positively with changes in transcript levels as measured by RNA-seq (Fig. 2c,d; see Methods). This suggests that the Spn1-dependent effects on transcript levels can be explained, at least in part, by changes in RNAPII occupancy. There was little effect of Spn1 depletion on either Ser5-P or Ser2-P levels, with Ser5-P retaining its expected preferential enrichment near transcription start sites and Ser2-P retaining its expected progressive enrichment over gene bodies^49^ (Fig. 2b, Supplementary Fig. 2f).

### 5’ regulatory regions influence Spn1-dependent regulation of transcript levels

Although Spn1 is thought of primarily as a transcription elongation factor, it was also shown to function at the promoter of the *S. cerevisiae CYC1* gene^11,16^. Therefore, we wanted to determine whether 5’ regulatory regions, including promoters, might play a role in Spn1-dependent regulation of transcript levels. To test this possibility, we made gene fusions of the 5’ regulatory regions of *UBI4* or *GCV3*—two genes respectively upregulated or downregulated in Spn1-depleted cells (Fig. 2d, Supplementary Fig. 2b)—to the coding region of *YLR454W,* a gene whose transcript levels were minimally affected by Spn1 depletion (Fig. 3a). The sequence used as the 5’ regulatory region of a gene included the 5’ untranslated region (UTR) of the gene, in addition to a promoter region defined using TSS-seq and TFIIB ChIP-nexus data (see Methods). These fusions and controls were constructed in a Spn1 degron strain.

**Figure 3.**
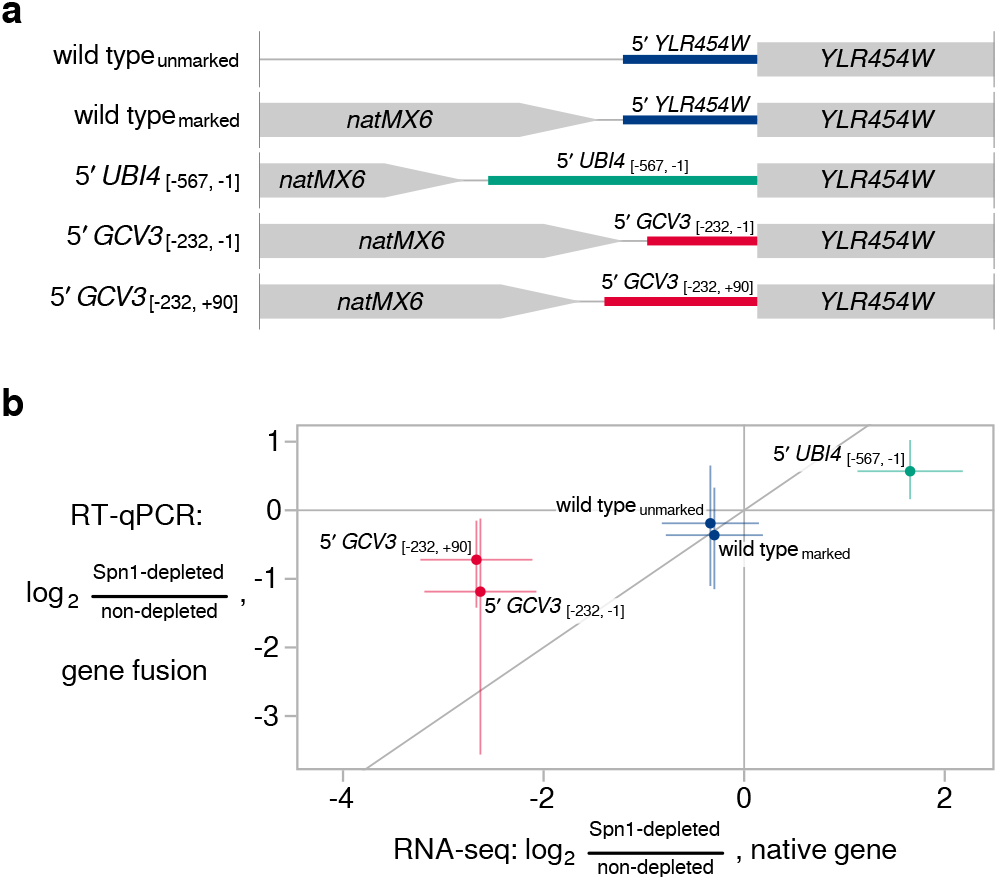
5*′* regulatory regions influence Spn1-dependent regulation of transcript levels. **a** Schematic of gene fusions in which the 5*′* regulatory regions of two Spn1-dependent genes replaced the 5*′* regulatory region of the YLR454W gene. The nucleotide coordinates of the native UBI4 and GCV3 regulatory regions are shown, relative to the start of the coding sequence^19,81^. All fusions were made in the Spn1 degron background. **b** Scatterplot summarizing results of the 5*′* regulatory region replacement experiment. The y-axis shows RT-qPCR measurements of the change in YLR454W transcript abundance upon Spn1 depletion in the gene fusion strains. The x-axis shows RNA-seq measurements of the change in transcript abundance upon Spn1 depletion for the native gene whose promoter was used to drive YLR454W expression in the fusion strains. The line y=x is drawn for comparison. Error bars indicate 95% confidence intervals, and points corresponding to the same native gene are slightly jittered by adding a small amount of random noise along the x-axis for clarity.

We then used RT-qPCR to measure the change in *YLR454W* transcript levels upon Spn1 depletion in the fusion and control strains (Fig. 3b). For the 5′*UBI4-YLR454W* fusion, transcript levels increased in response to Spn1 depletion about 48% relative to the non-depleted condition. For the 5′*GCV3-YLR454W* fusions, transcript levels decreased about 56% or 39%, depending on the particular sequence used for the *GCV3* 5’ regulatory region (see Methods). The direction of these changes is consistent with the RNA-seq and RT-qPCR data for the native *UBI4* and *GCV3* genes, although the magnitude of the *YLR454W* transcript changes in the fusions was smaller. Therefore, our results suggest that the 5’ regulatory regions of these genes contribute, at least partially, to the Spn1 dependence of their transcript levels.

### The association of Spn1 with RNAPII is dependent on Spt6

Spn1 directly interacts with Spt6, and both proteins are part of the RNAPII elongation complex^8–10,14,17,18^. To assess the possible dependency of one factor on the other for association with RNAPII, we performed immunopurification of the epitope-tagged RNAPII subunit Rpb3 under conditions where we depleted either Spn1 or Spt6. In the Spn1-depleted condition, Spt6 co-immunoprecipitated with Rpb3 at levels similar to the non-depleted condition (Fig. 4a, compare lanes 13 and 14, Spt6 panel). Therefore, the association of Spt6 with RNAPII is independent of Spn1, consistent with the fact that Spt6 directly interacts with RNAPII^50,51^. In contrast, in Spt6-depleted conditions, the interaction of Spn1 with RNAPII was lost (Fig. 4a, compare lanes 17 and 18, Spn1 panel). Therefore, the association of Spn1 with RNAPII is Spt6-dependent, supporting previous findings from studies of Spn1 (Iws1) in human cell lines^13,14^.

**Figure 4.**
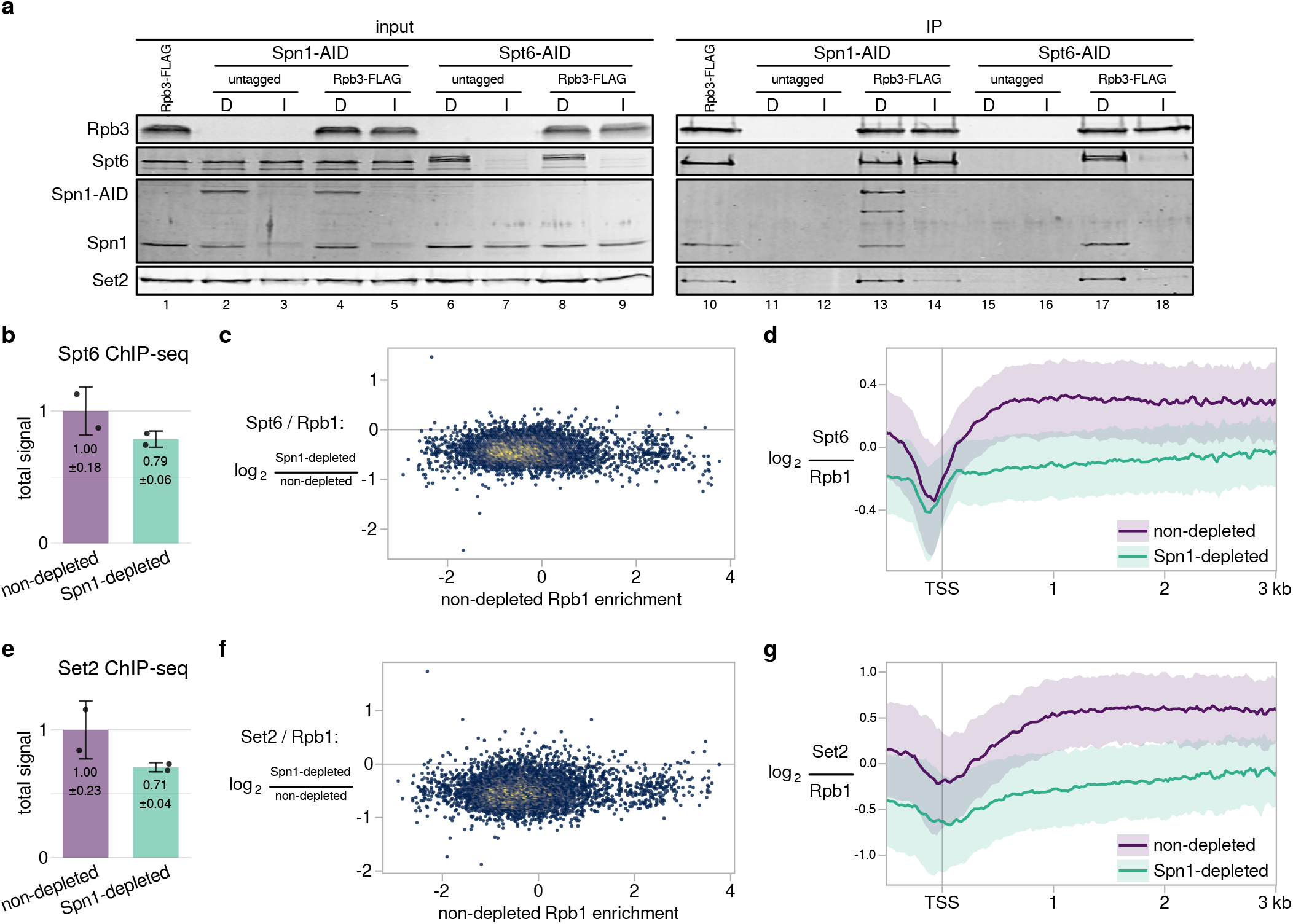
Requirement of Spn1 for Spt6 and Set2 association with RNAPII and chromatin. **a** Western blots showing the results of co-immunoprecipitation experiments to analyze the effects of Spn1 or Spt6 depletion on the binding of Spt6, Spn1, and Set2 to Rpb3-FLAG. Spn1 or Spt6 degron strains (Spn1-AID or Spt6-AID) were treated for 90 minutes with DMSO (lanes marked ‘D’) or 25 μM IAA (lanes marked ‘I’), after which Rpb3-FLAG was immunoprecipitated and co-immunoprecipitation of Spn1, Spt6, and Set2 was assayed by western blots using native antibodies. Rpb3-FLAG levels were measured using anti-FLAG antibody, and an Rpb3-FLAG strain lacking the degron tag on Spn1 or Spt6 was included as an additional non-depleted control. Shown is one representative western blot of three biological replicates with similar results. **b** Barplot showing total levels of Spt6 on chromatin in non-depleted and Spn1-depleted conditions, estimated from ChIP-seq spike-in normalization factors (see Methods). The values and error bars for each condition indicate the mean ± standard deviation of two replicates. **c** Scatterplot showing changes in Rpb1-normalized Spt6 ChIP enrichment upon Spn1 depletion versus non-depleted Rpb1 enrichment for 5,091 verified coding transcripts. Rpb1 enrichment values are the relative log_2_ enrichment of IP over input. **d** Average Rpb1-normalized Spt6 ChIP enrichment over 3,087 non-overlapping, verified coding genes aligned by TSS in non-depleted and Spn1-depleted conditions. The solid line and shading are the median and interquartile range of the ratio calculated from the mean of two Spt6 and four Rpb1 replicates. **e** Same as in (b) but for Set2. **f** Same as in (c) but for Set2. **g** Same as in (d) but for Set2.

Although our co-immunoprecipitation experiments indicated that Spn1 is not required for the association of Spt6 with RNAPII, it was possible that Spn1 was required for Spt6 recruitment at certain locations in the genome. For example, previous studies showed that Spn1 is required for the recruitment of Spt6 upon induction of the *CYC1* gene^11^. Therefore, to investigate the Spn1-dependence of Spt6 recruitment to chromatin genome-wide, we performed Spt6 ChIP-seq using a Spn1 degron strain expressing epitope-tagged Spt6. In the non-depleted condition, Spt6 and Rpb1 occupancy correlated strongly, as expected^10,20,52^ (Supplementary Fig. 3a), with the ratio of Spt6 to Rpb1 decreasing slightly with increasing Rpb1 occupancy (Supplementary Fig. 3b). Like in the case of Spn1 (Supplementary Fig. 1c), this suggests that Spt6 does not increase in equal proportion to RNAPII on genes as a function of transcription frequency. Following Spn1 depletion, Spt6 protein levels were unaffected (Supplementary Fig. 3c), while the total amount of Spt6 on chromatin was reduced to approximately 79% of non-depleted levels (Fig. 4b). The decrease in Spt6 on chromatin occurred uniformly over genes, with a net effect of reducing the ratio of Spt6 to Rpb1 over gene bodies (Fig. 4c,d). These results show that Spn1 is required for the optimal recruitment of Spt6 to chromatin.

### The association of Set2 with RNAPII and chromatin is dependent on Spn1

As both Spt6 and Spn1 are involved in the recruitment and/or activity of the histone methyltransferase Set2^5,14,24,25,27,28^, we also measured the association of Set2 with Rpb3 by co-immunoprecipitation in both Spn1-depleted and Spt6-depleted conditions. Our results showed that both proteins are required for Set2 association with RNAPII (Fig. 4a, compare lane 13 to 14 and lane 17 to 18, Set2 panel). In the case of Spt6 depletion, the diminished interaction of Set2 with Rpb3 can likely be explained by both the effect on Spn1 and the decreased Set2 protein levels that occur when Spt6 is mutant or depleted, as previously observed by others^27,28^, although this effect was modest in our results here (Fig. 4a, compare lane 6 to 7 and lane 8 to 9, Set2 panel).

To examine the Spn1-dependence of Set2 recruitment to chromatin across the genome, we performed Set2 ChIP-seq using a Spn1 degron strain expressing epitope-tagged Set2. In the non-depleted condition, we observed that the level of Set2 over coding genes positively correlated with Rpb1 levels (Supplementary Fig. 3a), while the ratio of Set2 to Rpb1 strongly negatively correlated with Rpb1 levels (Supplementary Fig. 3b). This suggests that the level of Set2 over genes varies much less with the level of transcription compared to elongation factors like Spt6 and Spn1 (Supplementary Figs. 3c and 1c). Following Spn1 depletion, the total amount of Set2 on chromatin was reduced to approximately 71% of non-depleted levels (Fig. 4e), and the ratio of Set2 to Rpb1 was reduced over gene bodies (Fig. 4f,g). The reduction in Set2 recruitment to chromatin measured by ChIP-seq appeared to be less severe than the reduction in Set2-RNAPII association observed by co-immunoprecipitation. This may be accounted for by technical differences between the two assays, such as the use of crosslinking in ChIP-seq, and may reflect the stabilization of Set2 on chromatin by factors besides RNAPII^27,28^, including other elongation factors^53,54^, nucleosomes/DNA^55–57^, and mRNA^58^. Thus, we conclude that Spn1 is required for the optimal recruitment of Set2 to chromatin, primarily by maintaining the Set2-RNAPII interaction.

### H3 levels are reduced near the 5’ end of a small set of genes after Spn1 depletion

As Spn1 binds to histones H3/H4 *in vitro*^16^ and has a modest role in the recruitment of Spt6 to chromatin (Fig. 4b-d), we investigated a possible role for Spn1 in the control of histone occupancy and distribution *in vivo.* To do this, we performed ChIP-seq for histone H3 in Spn1-depleted and non-depleted conditions. We observed that the effect of Spn1 depletion on H3 occupancy over coding genes varied with expression level: while most genes had minimally affected H3 occupancy and distribution after Spn1 depletion, a small set of 77 mostly highly expressed genes had significantly decreased H3 occupancy (Supplementary Fig. 4a, Fig. 5a). Furthermore, the decrease in H3 occupancy at these genes occurred specifically over the first ~500 bp of the gene (Fig. 5b,c). Since a 5’-specific effect might be masked for long genes when quantifying signal over the entire gene body, we then quantified the change in H3 occupancy over the first 500 bp of genes. By this approach, we observed the same relationship between expression level and H3 occupancy change and identified a total of 134 genes with significantly decreased H3 occupancy near the 5’ end (Supplementary Fig. 4b).

**Figure 5.**
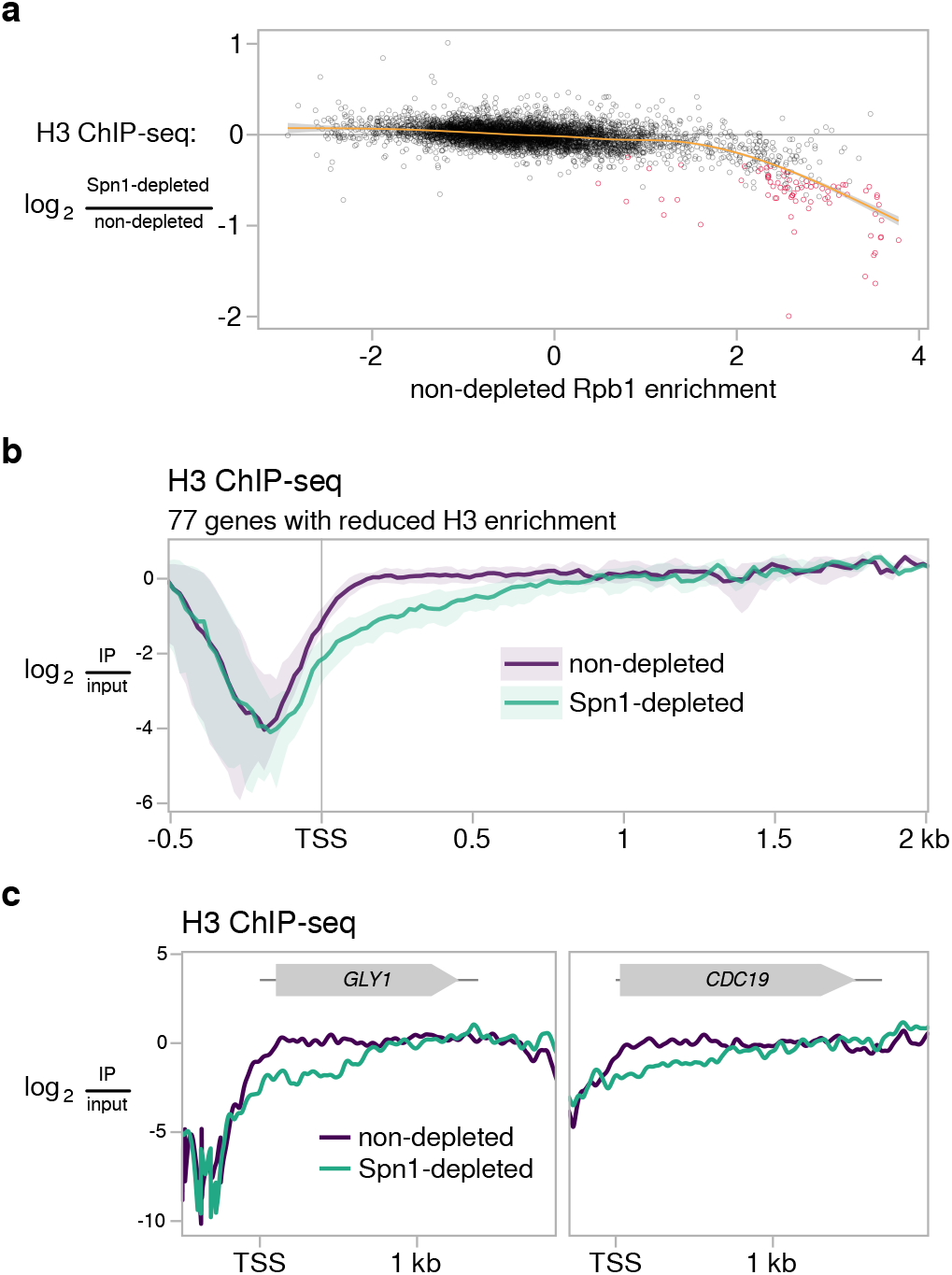
Histone H3 levels are reduced at the 5*′* end of a small set of genes after Spn1 depletion. **a** Scatterplot showing change in H3 enrichment upon Spn1 depletion versus non-depleted Rpb1 enrichment for 5,091 verified coding genes. Rpb1 enrichment values are the relative log_2_ enrichment of IP over input. Genes with significant decreases in H3 enrichment are colored red, and a cubic regression spline is overlaid. **b** Average H3 ChIP enrichment in non-depleted and Spn1-depleted conditions for the 77 genes with significantly decreased H3 ChIP enrichment over the entire gene upon Spn1 depletion. The solid line and shading are the median and interquartile range of the mean enrichment over four replicates. **c** H3 ChIP enrichment in non-depleted and Spn1-depleted conditions for two genes with significantly decreased H3 ChIP enrichment upon Spn1 depletion. The mean enrichment over four replicates is shown.

The 5’ reduction in H3 at highly expressed genes upon Spn1 depletion was reminiscent of the 5’ reduction in H3 over a broader set of genes previously observed upon Spt6 depletion^59^ and in an *spt6* mutant with decreased Spt6 protein stability^23^. Because of this and the known histone chaperone function of Spt6, we examined whether the reduction in H3 after Spn1 depletion could be explained by reduced levels of Spt6 on chromatin (Fig. 4b-d). We found that for those genes with reduced H3 levels, the H3 changes did not correlate with changes in Rpb1-normalized Spt6 signal (Supplementary Fig. 4c). Additionally, the 5’ reduction in H3 at those genes was coincident with a 5’ reduction in Rpb1 (Supplementary Fig. 4d). These results suggest that the changes in H3 occupancy at this set of genes upon Spn1 depletion are not the indirect effect of changes in Spt6 recruitment and may be related to changes in transcription, though we cannot say if the relationship is causal.

### Spn1 is required for the normal distribution of co-transcriptional histone modifications

Given previous studies that suggested a role for Spn1 in H3K36 methylation^3,5,14^ and the change we observed in Set2 recruitment after Spn1 depletion (Fig. 4e-g), we used ChIP-seq to measure the genome-wide occupancy of H3K36me2 and H3K36me3 in Spn1-depleted and non-depleted conditions. In these experiments, we also included analysis of H3K4 trimethylation (H3K4me3), given the genetic relationship between Spn1 and the enzymes required for the methylation and demethylation of H3K4^21,22^.

For H3K36 methylation, our results showed that, while the total level of each histone modification on chromatin was not greatly affected by Spn1 depletion (Supplementary Fig. 5a,b), the distribution of each modification was markedly changed (Fig. 6a,b, Supplementary Fig. 5c). In the non-depleted condition, the H3K36 methylation marks were distributed as expected: H3K36me2 and H3K36me3 were primarily enriched within gene bodies, with H3K36me3 reaching its maximum levels further downstream of the TSS than H3K36me2^40,60^ (Fig. 6a,b). In contrast, depletion of Spn1 resulted in prominent 3’ shifts in the distributions of both H3K36me2 and H3K36me3 over the vast majority of coding genes (Fig. 6a,b). More specifically, in the non-depleted condition, the median H3-normalized H3K36me2 signal reached 90% of its maximum value within approximately 450 bp of the TSS, while in the Spn1-depleted condition, 90% of the maximum value was not reached until about 650 bp (Fig. 6a, top). For H3-normalized H3K36me3 in the non-depleted condition, the median signal reached 90% of its maximum value within about 690 bp of the TSS, while in the Spn1-depleted condition, 90% of the maximum value was not reached until about 1390 bp (Fig. 6b, top). Though these changes in distribution occurred at most genes, the magnitude of the changes increased with increasing expression level (Supplementary Fig. 5d).

**Figure 6.**
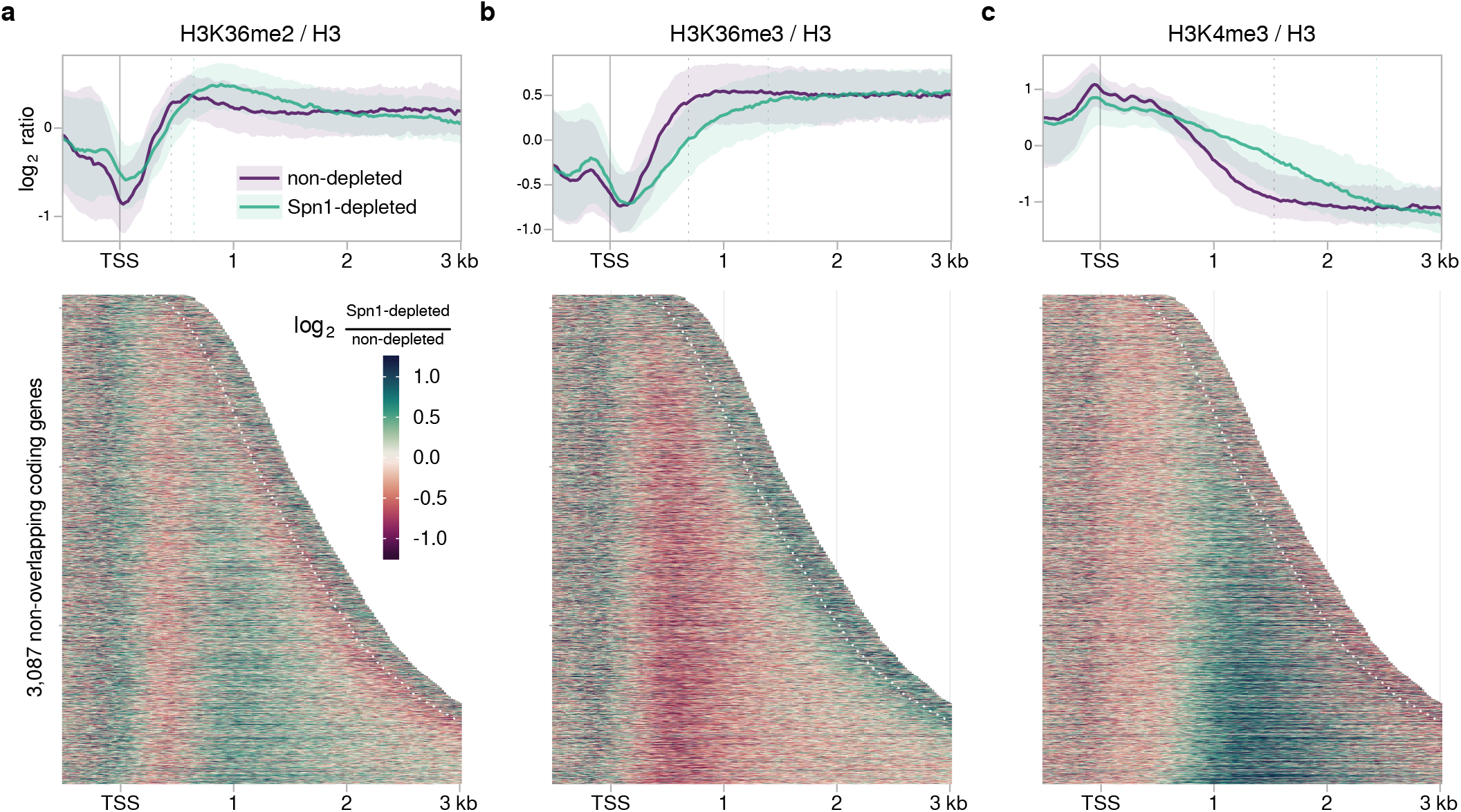
Spn1 is required for normal localization of histone H3 modifications along coding genes. **a** Top: Average H3-normalized H3K36me2 for 3,087 non-overlapping, verified coding genes aligned by TSS in non-depleted and Spn1-depleted conditions. The solid line and shading are the median and interquartile range of the mean ratio over two replicates. Vertical dotted lines indicate the distances at which the median signal in each condition reached 90% of its maximum value. Bottom: Heatmap of the change in H3-normalized H3K36me2 enrichment upon Spn1 depletion for the same genes, aligned by TSS and arranged by transcript length. Data is shown for each gene up to 300 bp 3*′* of the cleavage and polyadenylation site, which is indicated by the white dotted line. **b** Same as in (a) but for H3K36me3. **c** Same as in (a) but for H3K4me3, with vertical dotted lines instead indicating the distances at which the median signal in each condition dropped to 10% of its maximum value.

The genome-wide distribution of H3K4me3 was also strongly affected by Spn1 depletion. While in the non-depleted condition, H3K4me3 was primarily localized over the 5’ regions of genes, as expected^38–40,61^, depletion of Spn1 caused H3K4me3 signal to spread toward the 3’ end for the vast majority of genes, into regions normally enriched for H3K4me2 and H3K4me1^40^ (Fig. 6c, Supplementary Fig. 5c). In the non-depleted condition, the median H3-normalized H3K4me3 signal over coding genes decreased to 10% of its maximum value within about 1530 bp of the TSS, whereas in the Spn1-depleted condition, the median signal did not decrease to 10% of its maximum value until about 2430 bp (Fig. 6c, top). As with H3K36 methylation, the magnitude of the change in H3K4me3 distribution over genes increased with increasing expression level (Supplementary Fig. 5d). It is possible that the change in H3K4me3 pattern upon Spn1 depletion is caused by altered recruitment or activity of either Set1, the H3K4 methyltransferase, or Jhd2, the demethylase, though this remains to be tested. Altogether, we conclude that Spn1 is required to maintain the normal pattern of H3K36 and H3K4 methylation across the genome.

### Spn1 depletion results in increased intron retention in ribosomal protein transcripts

. Since both Spn1 and H3K36 methylation have been previously implicated in splicing^3,5,62–66^, we looked for changes in splicing efficiency after Spn1 depletion by using our RNA-seq data to estimate intron retention (see Methods). We observed a general tendency for increased intron retention after Spn1 depletion (Fig. 7a), particularly in ribosomal protein transcripts, whose high expression levels gave us greater power to detect changes in intron retention. Out of 252 introns analyzed, we found 59 introns with significantly increased retention upon Spn1 depletion and two introns with significantly decreased retention (Fig. 7a). Of the introns with significantly increased retention after Spn1 depletion, 27/59 showed increases of at least 10%, and 54/59 were from ribosomal protein transcripts. To verify our intron retention estimates, we used RT-qPCR to measure the ratio of unspliced to spliced transcripts for four genes that showed increased intron retention and two genes without significant changes (Fig. 7b). In all cases, the changes in unspliced transcript abundance observed by this method were in agreement with the results from the transcriptome-wide analysis.

**Figure 7.**
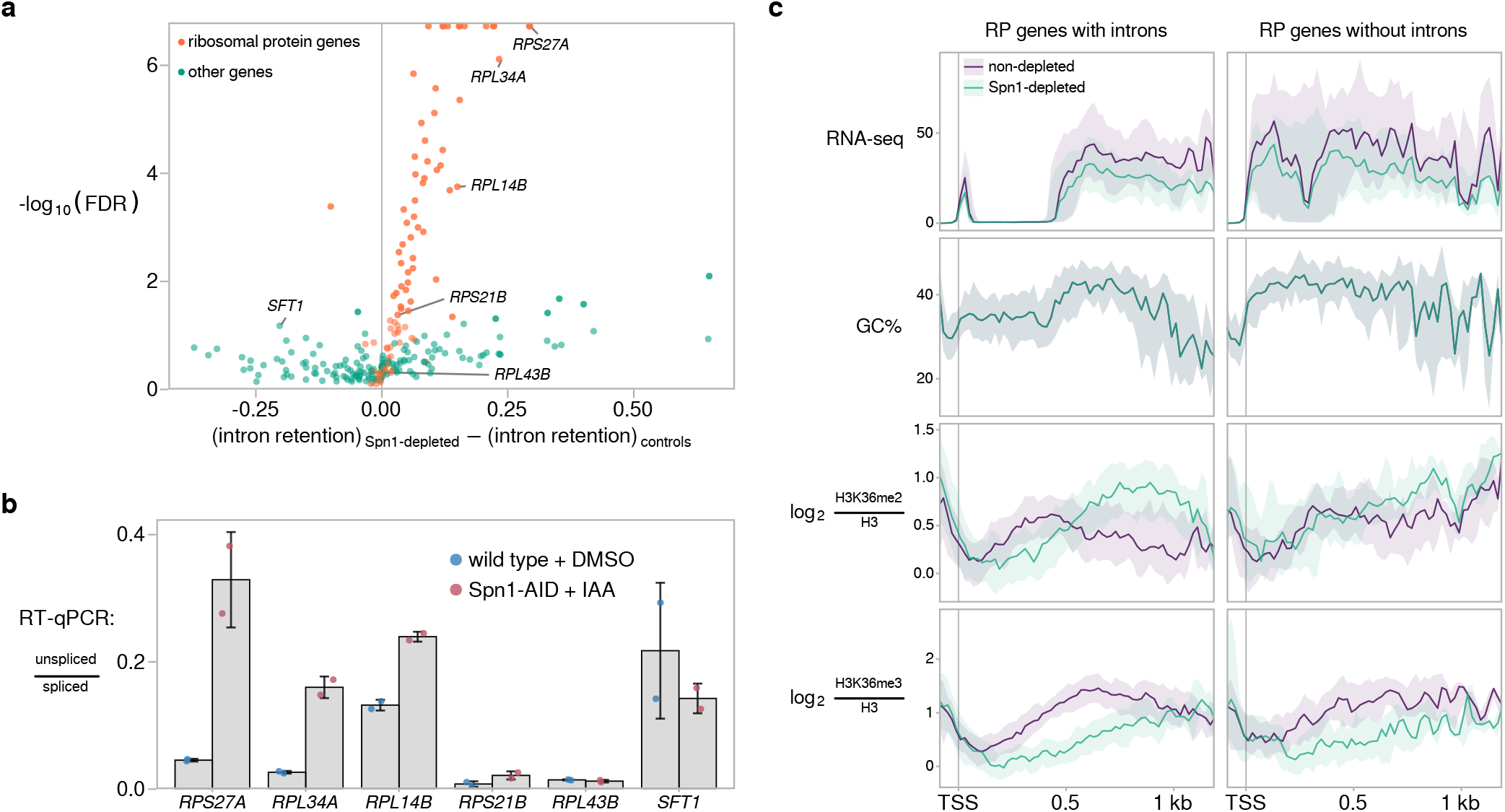
Depletion of Spn1 results in increased intron retention in ribosomal protein transcripts. **a** Volcano plot showing changes in intron retention upon Spn1 depletion for 252 introns encoded from within coding regions of nuclear genes. Intron retention is defined as the proportion of unspliced transcripts and was estimated from RNA-seq data (see Methods). Labeled introns were further assayed by RT-qPCR, as shown in (b). **b** RT-qPCR measurements of the ratio of unspliced to spliced transcripts in a wild-type strain treated with DMSO for 90 minutes and the Spn1 degron strain treated with 25 μM IAA for 90 minutes. Error bars indicate the mean ± standard deviation of two replicates. **c** Visualization of average changes in H3K36 methylation state at ribosomal protein genes with and without introns. For each group of genes, the following data are plotted: sense strand RNA-seq coverage, GC% in a 21-bp window, and H3-normalized H3K36me2 and H3K36me3 ChIP enrichment. The solid line and shading are the median and interquartile range over the genes in each group. The introns of RP genes tend to be close to the 5’ end of the gene, the region where RNA-seq signal is depleted (top left panel).

H3K36 methylation has been linked to splicing in mammalian cells by studies showing that chromodomain and PWWP domain-containing proteins such as MRG15 can bind SETD2-dependent H3K36me3 to bridge the association of splicing factors and chromatin, thus co-transcriptionally regulating alternative splicing^62,65,66^. Eaf3, the yeast ortholog of MRG15, also binds to both H3K36me3 and H3K36me2^24,33,34,67^ and mediates the recruitment of splicing factors^64^ supporting previous results that implicated H3K36 di- and/or trimethylation in splicing regulation in yeast^63^. Therefore, we wanted to determine if there were Spn1-dependent changes in either H3K36me2 or H3K36me3 that were specific to intron-containing genes. Sixty-five percent of the 137 ribosomal protein (RP) genes contain introns, giving us an opportunity to compare two sets of genes similar in expression and length but differing by the presence or absence of introns. In the non-depleted condition, intron-containing RP genes showed a slightly higher level of H3K36me2 over the intron compared to the downstream exon (Fig. 7c). Following Spn1 depletion, redistribution of this modification toward the 3’ end resulted in a marked decrease in signal over the intron, a pattern not observed at a similar distance from the TSS in RP genes without introns. Additionally, RP genes with introns had a slightly greater decrease in H3K36me3 over most of the gene body compared to RP genes without introns. We did not detect major differences in the distribution of other factors or modifications assayed between the two sets of genes, apart from a very mild increase in Rpb1 Ser5-P over the introns, which has been observed when pre-spliceosome formation is perturbed^68,69^ (Supplementary Fig. 6). We obtained very similar results when we performed this analysis comparing intron-containing RP genes to non-RP genes without introns matched by length and expression (data not shown). Finally, the overlap of decreases in H3K36me2 and regions encoding introns observed for RP genes was not seen for non-RP intron-containing genes, as the decreases in H3K36me2 over non-RP intron-containing genes occurred further 3’ of the TSS and 5’ splice site (Supplemental Fig. 6b). Thus, for intron-containing RP genes, distinct changes in H3K36me2 and H3K36me3 correlate with increased intron retention.

## Discussion

In this study, we have uncovered previously unknown *in vivo* roles for Spn1, an essential and conserved factor. We have shown that Spn1 is widely required for normal transcript levels, as depletion of Spn1 significantly reduces mRNA levels of over 1,500 genes in a manner that correlates with changes in Rpb1 occupancy. In addition, depletion of Spn1 alters the distribution of H3K36 and H3K4 methylation along most genes and impairs efficient splicing, particularly in ribosomal protein transcripts. Furthermore, Spn1 is required for normal levels of histone H3 over the 5’ ends of a small set of mostly highly transcribed genes, providing the first genome-wide, *in vivo* evidence supporting recent *in vitro* studies that implicated Spn1 as a histone chaperone^16^. Finally, we have shown that the association of Spn1 with RNAPII is dependent on its binding partner, Spt6. Together, these results establish that Spn1 plays a genome-wide role in transcription and chromatin dynamics.

We have shown that a major function of Spn1 is to maintain proper localization of H3K36me3, H3K36me2, and H3K4me3, as Spn1 depletion causes redistribution of these three transcription-coupled histone modifications toward the 3’ ends of genes. Mislocalization of these histone marks has been previously observed in other transcription and chromatin mutants in both yeast and mammalian cells, although in some cases the shifts were in the 5’ direction, opposite to the shifts observed after Spn1 depletion. For example, loss of the chromatin remodeler Chd1 resulted in 5’ shifts of both H3K36me3 and H3K4me3^70,71^. Interestingly, the loss of Chd1 also resulted in more efficient splicing of ribosomal protein transcripts, in contrast to the less efficient splicing that we observed after Spn1 depletion. Thus, Spn1 and Chd1 appear to play opposite roles. In the case of Spn1 depletion, it is plausible that reduced Set2 recruitment and/or activity contribute to the changes in H3K36 methylation pattern.

Changes in the rate of transcription have also been shown to cause shifts of H3K36 and H3K4 methylation. In yeast, an RNAPII slow mutant with a reduced polymerization rate showed a 5’ shift of H3K4me2 and H3K4me3, while an RNAPII fast mutant with an increased polymerization rate showed a 3’ shift of these modifications^61,72^. In mammalian cells, a 5’ shift of H3K36me3 was also observed in an RNAPII mutant with slowed transcription rate^73^, opposite to what we observed after Spn1 depletion. In both the yeast and mammalian studies, the histone modification changes correlated with shifts of Rpb1 Ser2-P: a 3’ shift in slow mutants and a 5’ shift in the fast mutant^61,73^. In contrast, we did not detect a significant change in Rpb1 Ser2-P after Spn1 depletion; therefore, the histone modification 3’ shifts that we observe might not be caused by an increased rate of transcription, although this remains to be tested.

A recent study showed that the histone chaperones Spt6 and FACT (composed of Spt16 and Pob3) are also required for the normal distribution of several histone modifications, as shifts toward the 3’ end occurred in *spt6* and *spt16* mutants^72^. Evidence suggested that these shifts were a consequence of transcription-dependent loss of histones on chromatin due to impaired nucleosome reassembly in the wake of transcription. The decreases in histone H3 occupancy that we observed after Spn1 depletion were much less extensive than those observed in the *spt6* and *spt16* mutants. Therefore, although Spn1 is likely required for nucleosome reassembly at highly expressed genes, a defect in nucleosome assembly is unlikely to completely explain the widespread shifts in histone modifications observed after Spn1 depletion.

We found that Spn1 depletion causes both increased retention of many ribosomal protein introns and reduced H3K36me2 and H3K36me3 over the regions encoding these introns. The possibility that the reduced H3K36 methylation causes the defects in splicing is supported by previous studies showing that loss of H3K36 methylation impairs splicing from yeast to mammals^5,62–66,70,74^. We observed that Spn1 depletion caused ribosomal protein genes to have reduced H3 and H3K36me3 regardless of whether the gene contained an intron; however, ribosomal protein genes with introns had a distinct redistribution of H3K36me2 after Spn1 depletion, resulting in decreased levels of this mark before the 3’ splice site and increased levels thereafter. Our results, then, suggest that H3K36me2 is a Spn1-dependent mark that distinguishes ribosomal protein genes with introns from those without. Given that defects in splicing have also been observed to alter the recruitment of SETD2 and the distribution of H3K36me3 at intron-containing genes in mammalian cells^75,76^, further experiments will be needed to determine whether the direct effect of Spn1 depletion is on H3K36 methylation patterns or splicing.

Our studies have also revealed new insights into the molecular relationship between Spn1 and Spt6. The importance of this interaction is highlighted by the fact that *spn1* or *spt6* mutations that impair this interaction cause severe mutant phenotypes^17,18^. Other studies previously suggested that the two proteins do not contribute to a common essential function since they differ based on both genetic and molecular properties^10,11,18^. Our results extend these distinctions, as depletion of Spn1 causes different effects on transcription compared to mutation or depletion of Spt6, particularly with respect to the level of antisense transcription^19,26,48^. The critical nature of the Spn1-Spt6 interaction can be explained by our findings that the association of Spn1 with RNAPII is strongly dependent on Spt6, and that Spn1 is required for optimal recruitment of Spt6 to chromatin.

In summary, our studies have demonstrated that Spn1 is widely required *in vivo* for normal transcript levels, splicing, and chromatin-related functions. Given its physical and genetic interactions with two other essential histone chaperones, Spt6 and FACT^21,22,77^, and its genetic interactions with additional histone chaperones, including Asf1 and the Hir complex^16^, it appears that Spn1 is part of a network of histone chaperones required for normal gene expression. Deciphering the individual roles of these factors and how they function together remains a challenge for future studies.

## Methods

### Yeast strains

All yeast strains are listed in Supplementary Table 1, and all primers used for strain constructions are listed in Supplementary Table 2. *S. cerevisiae* liquid cultures were grown in YPD (1% yeast extract, 2% peptone and 2% glucose) at 30°C. *S. pombe* liquid cultures were grown in YES (0.5% yeast extract, 3% glucose, 225 mg/L each of adenine, histidine, leucine, uracil, and lysine) at 32°C. For Spn1 and Spt6 depletion experiments using an auxin-inducible degron (AID)^78^, strains expressing the *Oryza sativa TIR1* gene and *SPN1-AID* or *SPT6-AID* were constructed as previously described^79^. For Spt6 ChIP-seq, a 5’ HA-tagged allele of *SPT6* was crossed into the *SPN1-AID* background. For Set2 ChIP-seq, Set2 was epitope-tagged by transformation of a DNA fragment amplified from pFA6a-3xHA-NatMX6 (FB2517)^80^, which fuses the HA epitope to the C-terminal end of Set2. For co-immunoprecipitation experiments, a triple Flag-tagged allele of *RPB3* was crossed into the *SPN1-AID* and *SPT6-AID* backgrounds. In all cases, possible growth effects of epitope tags were assessed by colony size. For 5’ regulatory region replacement experiments, the *YLR454W* promoter and 5’ UTR were replaced by those of *UBI4* or *GCV3*. To do this, the *UBI4* and *GCV3* 5’ regulatory regions were amplified from genomic DNA of a wild-type strain (FY92) using a forward primer with a 5’ 20-bp overhang that would allow for later amplification of the NatMX cassette from a pFA6a plasmid (this 20-bp sequence is the reverse complement of the reverse primer sequence used for making deletions in^80^). The reverse primer for 5’ regulatory region amplification included a 3’ overhang with homology to the 40-bp sequence beginning at the *YLR454W* start codon. The resulting amplicon containing homology to the pFA6a-NatMX plasmid, the regulatory region of interest, and homology to the region beginning at the *YLR454W* ATG was recovered using the E.Z.N.A. Cycle Pure Kit and used as a reverse primer in a second PCR reaction in which the NatMX cassette was amplified from the pFA6a plasmid. The forward primer for this reaction included 40 bp of homology to the region from −323 to −284 relative to the *YLR454W* start codon and the Longtine forward primer sequence^80^ for making deletion cassettes from pFA6a. The reverse primer for the first PCR reaction was used additionally in this second PCR to increase product yield. The final PCR product for each construct was ethanol-precipitated, gel-purified using the E.Z.N.A. Gel Purification Kit, and used to transform *S. cerevisiae* strain FY3227 (Spn1-AID). Transformants were selected on YPD plates that contained 100 μg/mLnourseothricin, and insertion of each cassette in the correct orientation and location was checked by two PCR reactions (PCR-1 using NR179 and FO5520, PCR-2 using FO7627 and NR180). The PCR-2 reaction amplified a region from the NatMX cassette to the first 379 bp of *YLR454W* ORF; this product was used to sequence across the 5’ regulatory region of each construct using primer FO6606. A control strain in which the NatMX cassette alone was inserted upstream of the endogenous *YLR454W* promoter without deleting any *YLR454W* sequences was also constructed. In this case, only one PCR reaction using the pFA6a-NatMX plasmid as a template was required. The *YLR454W* 5’ regulatory region was defined as the region from −283 to −1 relative to the ATG and included the TFIIB (−208 to −103 relative to the ATG) and TSS (−148 to −90 relative to the ATG) peaks previously identified^19^. The *UBI4* 5’ regulatory region was defined as the region between −567 and −1 relative to the ATG and included the TFIIB (−220 to −80 relative to the ATG) and TSS (−111 to −87 relative to the ATG) peaks, as well as other regulatory elements previously described^81^. Two different 5’ regulatory region amplicons were made for *GCV3*: one in which it was defined as the region between −232 and −1 relative to the annotated ATG and one in which it was defined as the region between −232 and +90 relative to the ATG (this ATG remained in frame with the *YLR454W* ATG in the final strain). The latter construct was made because the TFIIB peak mapped in Doris et al. spanned ~40 bp into coding region of *GCV3* (TFIIB peak: −114 to +43 relative to the ATG; TSS peak: −15 to −1 relative to the ATG). In summary, the *UBI4* and *GCV3* 5’ regulatory region amplicons, marked with NatMX upstream, were inserted immediately upstream of the *YLR454W* start codon, while the region from −283 to −1 was deleted in the process.

### Auxin-induced depletion of Spn1 and Spt6

Spn1 depletion was optimized as previously described^79^ by testing final concentrations of the auxin 3-indoleacetic acid (IAA), ranging from 25-500 μM and treatment times ranging from 0-120 minutes. Viability was measured by counting colony-forming units per OD_600_ unit plated in the 25-μM IAA time-course. Optimization of the time for Spt6 depletion was done for the 25-μM IAA treatment. For all Spn1 and Spt6 depletion experiments in this study, IAA was added to a final concentration of 25 μM followed by growth for 90 minutes.

### Western blotting and antibodies

For checking Spn1 and Spt6 protein levels in all depletion experiments, OD-normalized aliquots of the cultures grown for the main experiment were saved and processed later on. For assessing levels of proteins and histone modifications assayed in ChIP-seq experiments, triplicate 20-mL cultures for the wild type, Spn1-noAID (contains TIR1 but no AID tag on Spn1), and Spn1-AID strains were grown to OD_600_ ≈ 0.7 and treated with DMSO or IAA as described in the ChIP-seq methods. Culture volumes were normalized by their OD_600_ to an OD equivalent (OD_600_ * volume of the culture) of 12 for harvesting. Protein extracts were prepared as described previously^27^, and 12-15 μL were loaded on 10% or 15% SDS-PAGE gels for western blotting. The following dilutions of primary antibodies were used: 1:1,000 anti-Rpb1 (Millipore, 8WG16); 1:6,000 anti-Spn1 (generously provided by Laurie Stargell); 1:10,000 anti-Spt6 (generously provided by Tim Formosa); 1:6,000 anti-Set2 (generously provided by Brian Strahl); 1:1,000 anti-H3K36me3 (Abcam, ab9050); 1:1,000 anti-H3K36me2 (Abcam, ab9049); 1:2,000 anti-H3 (Abcam, ab1791); 1:8,000 anti-Pgk1 (Life Technologies, #459250). The following dilutions of secondary antibodies were used: 1:10,000 goat anti-rabbit IgG (Licor IRDye 680RD) and 1:20,000 goat anti-mouse IgG (Licor IRDye 800CW). Quantification of western blots was done using ImageJ 1.48v.

### RNA-seq

Duplicate 100-mL cultures for the wild type (FY3225), Spn1-noAID (FY3226), and Spn1-AID (FY3227) strains were grown in YPD at 30°C to ~ 3.5 × 10^7^ cells/mL, diluted back to OD_600_ ≈ 0.3 in pre-warmed YPD, divided into two equal volumes, treated with DMSO or IAA at a final concentration of 25 μM, and grown for an additional 90 minutes. An OD-normalized aliquot of each culture was processed to verify depletion by western blot. Cells were counted and harvested, and each sample was spiked in to make a final concentration of 10% *S. pombe* cells and flash-frozen. Total RNA was prepared by hot acid phenol extraction^82^, and a portion of each sample was treated with RQ1 DNase and used for subsequent steps (Promega, M6101). rRNA was removed using a Ribo-Zero kit for yeast (Illumina, MRZY1324) as per manufacturer instructions, except all volumes were halved. Library preparation was carried out according to a previously published method, except fragmentation was performed prior to linker ligation for RNA-seq^83^. Briefly, alkaline fragmentation (2x fragmentation buffer: 100 mM NaCO_3_ pH 9.2, 2 mM EDTA pH 8.0) was performed to obtain 40-70-nucleotide fragments that were subsequently size-selected and dephosphorylated with T4 PNK (NEB, M0201S) for ligation of a 3’ DNA linker containing a random hexamer molecular barcode to allow detection of PCR duplicates^84^. First-strand cDNA was synthesized with SuperScriptIII (Invitrogen, #56575) using a primer with sequence complimentary to the 3’ linker and with an additional adapter sequence. cDNA was circularized with CircLigase (Epicentre, CL4111K). DNA libraries were amplified by PCR for 6-10 cycles with Phusion HF DNA Polymearse (NEB, M0530) using Illumina TruSeq indexing primers for multiplexing. Libraries were gel-purified and run on Agilent Bioanalyzer and/or TapeStation to assess size distribution, quality, and concentration. RNA-seq libraries were sequenced on the Illumina NextSeq 500 platform at the Biopolymers Facility at Harvard Medical School.

### RT-qPCR

Spn1-AID cells were cultured and treated with DMSO or IAA as described for RNA-seq. Total RNA was prepared by hot acid phenol extraction, DNase-treated, and ethanol-precipitated after each step. cDNA was synthesized from 1 μg of RNA using qScript XLT cDNA SuperMix (Quanta Biosciences, #95161), which contains a blend of oligo d(T) and random primers. A no-RT (no reverse transcription) control, where no cDNA SuperMix was added to the RNA, was included for each sample and used for qPCR. The 20-μL RT reactions were brought up to 100 μL with water, and three 1:3 serial dilutions were prepared for each sample. Brilliant III Ultra-Fast SYBR Green QPCR Master Mix (Agilent Technologies, #600882) was used for quantitative PCR. Primers are listed in Supplementary Table 1. Triplicate intercept values for regions of interest were divided by triplicate intercept values of *S. pombe pma1^+^* (primers FO7083 and FO7084) if *S. pombe* cells were used as spike-in. For intron retention analysis, unspliced transcripts were measured with a forward primer spanning the 3’ intron-exon junction, while spliced transcripts were measured with a forward primer spanning the exon-exon junction. The reverse primer was the same for both transcript species. The ratio of unspliced to spliced transcripts was calculated for six genes in duplicate. For the 5’ regulatory region replacement experiments, changes in *YLR454W* transcript levels following Spn1 depletion were measured using primers NR183 and NR184, which amplified the region from +994 to +1091 relative to the start codon.

### ChIP-seq

The Spn1-AID strain (FY3227) was used for ChIP of Rpb1, Rpb1 CTD modifications, histone H3, histone modifications, and Spn1 (Spn1-AID-3xV5). Spn1-AID strains containing HA-tagged Spt6 (FY3228) or Set2 (FY3229) were used for those respective ChIPs. For *S. cerevisiae* strains, duplicate cultures were grown in YPD at 30°C to OD_600_ ≈ 0.65, diluted back to OD_600_ ≈ 0.3-0.4 in pre-warmed YPD, divided into two equal volumes, treated with DMSO or IAA at a final concentration of 25 μM, and allowed to grow for 90 minutes. An OD-normalized aliquot of each culture was processed to verify depletion by western blot. Cultures were processed as previously described^27^, and the final pellet after cell lysis was resuspended in 500 μL LB140 and sonicated in a QSonica Q800R machine for 15 minutes (30 seconds on, 30 seconds off, 70% amplitude) to achieve 100-500-bp DNA fragments. For *S. pombe* strains, cultures were grown at 32°C in YES to OD_600_ ≈ 0.7-0.8 and processed similarly to the *S. cerevisiae* cultures, except for the following steps. Bead-beating was done for 10-14 minutes, chromatin was concentrated at the last resuspension in LB140, and samples were sonicated for 20 minutes. Protein concentrations of all *S. cerevisiae* and *S. pombe* chromatin samples were measured by Bradford^85^. For Spt6 and Set2 ChIPs, 1 mg of chromatin was used as input, whereas 500 μg of chromatin was used as input for all other ChIPs. As a spike-in control, *S. pombe* chromatin was added to a final concentration of 10%, and the volume was brought up to 800 μL with WB140 buffer. Samples for Rpb1, Spt6, Set2, H3, and H3K36me3 ChIP were spiked in with an *S. pombe* strain encoding an HA-tagged protein (FWP568). Samples for Rpb1, Rpb1 Ser5-P, Rpb1 Ser2-P, H3, H3K36me2, H3K4me2, and Spn1 were spiked in with an *S. pombe* strain encoding a V5-tagged protein (FWP485). Before addition of antibody, 5% (40 μL) was set aside as input and stored for later processing, starting at the decrosslinking step later on. The following antibodies were used for IPs: 10 μg anti-HA (Abcam, ab9110), 10 μL anti-Rpb1 (Millipore, 8WG16), 8 μg anti-H3 (Abcam, ab1791), 6 μg anti-H3K36me3 (Abcam, ab9050), 6 μg anti-H3K36me2 (Abcam, ab9049), 6 μg anti-H3K4me3 (Abcam, ab8580), 10 μg anti-Ser5-P (Active Motif, 3E8), 10 μg anti-Ser2-P (Active Motif, 3E10), 8 μL anti-V5 (Invitrogen, R960-25). Immunoprecipitation reactions were incubated at 4°C with end-over-end rotation overnight (~16-18 hr). Protein G Sepharose beads (GE Healthcare, 4 Fast Flow) were washed with WB140, and a 50% slurry was subsequently made in WB140. Fifty-μL aliquots of slurry were transferred into new Eppendorf tubes, and the WB140 was removed. The immunoprecipitation reactions were added directly to the beads and incubated at 4°C with end-over-end rotation for 4 hr. Bead-washing and elution were done are previously described^27^. The saved inputs were thawed from storage, and 60 μL LB140 and 100 μL TES were added. The 200-μL eluates and inputs were incubated at 65°C overnight to reverse crosslinking. RNase A/T1 and Proteinase K digestions, DNA purification, and library preparation were done as previously described^27^. The number of PCR cycles for library amplification was determined empirically for each sample and ranged from 11-19 cycles. Samples were again purified twice using 0.7x volume SPRI beads and run on Agilent Bioanalyzer to assess size distribution, quality, and concentration. ChIP-seq libraries were sequenced on the Illumina NextSeq 500 platform at the Harvard Bauer Core Facility.

### Co-immunoprecipitation

Rpb3-Flag (FY3224), Spn1-AID (FY3227), Rpb3-Flag Spn1-AID (FY3230), Spt6-AID (FY3122), and Rpb3-Flag Spt6-AID (FY3231) cells were cultured in triplicate and treated with DMSO or IAA as described for ChIP-seq. Depletions of Spn1 and Spt6, as well as levels of other proteins assayed in this experiment, were verified by western blot. After collecting the cell pellets and washing them with water, the cells were resuspended in an amount of IP buffer (20 mM HEPES pH 7.6, 20% glycerol, 1 mM DTT, 1 mM EDTA, 125 mM KoAc, 1% NP-40) proportional to the OD of the culture harvested (with a culture at OD_600_ 0.65 receiving 500 μL of buffer). The cells were lysed by bead-beating at 4°C for 6 minutes with 2-minute incubations on ice after every minute of bead-beating. The lysates were collected, centrifuged at 14,000 rpm for 10 minutes at 4°C, and the cleared extracts were transferred to new tubes. Protein concentrations of all extracts were measured using Pierce BCA Protein Assay Kit (Thermo Fisher, #23225). Samples for immunoprecipitation were prepared at a final concentration of 15 μg/μL protein in IP buffer up to 500 μL. As input, 40 μL of each sample was set aside, 20 μL 3x SDS buffer (240 mM Tris-HCl pH 6.8, 6% SDS, 16% β-mercaptoethanol, 4% SDS, 0.6 mg/mL bromophenol blue, 30% glycerol) was added, and this was frozen until ready to use. Anti-Flag M2 affinity gel beads (Sigma, A2220) were washed once in IP buffer and prepared as a 10% bead slurry in IP buffer. One hundred-μL aliquots of slurry (containing a total of 10 μL packed beads) were prepared in separate tubes, and 400 μL of the normalized extracts were added to each tube and incubated at 4°C with end-over-end rotation for 2 hr. Following the IP, the beads were washed three times with 500 μL of IP buffer for 2 minutes per wash at 4°C. For elution, the beads were resuspended in 45 μL of 3x SDS buffer and boiled for 5 minutes. The beads were then spun down, and the supernatant was transferred to a new tube and either stored or used for western blot. The following antibodies were used for western blot: 1:8,000 anti-Flag (Sigma, F3165); 1:6,000 anti-Spn1 (provided by Laurie Stargell); 1:10,000 anti-Spt6 (provided by Tim Formosa); 1:6,000 anti-Set2 (provided by Brian Strahl); 1:2,000 anti-V5 (Invitrogen, R960-25) to check depletions.

### Data management

All data analyses were managed using the Snakemake workflow management system^86^. An archive containing code and raw data suitable for reproducing the analyses presented in this manuscript is available at Zenodo (doi.org/10.5281/zenodo.3901642). Additionally, updated versions of the Snakemake pipelines used are available at github.com/winston-lab.

### Genome builds and annotations

The genome builds used were *S. cerevisiae* R64-2-1^87^ and *S. pombe* ASM294v2^88^. *S. cerevisiae* transcript coordinates were generated from TIF-seq^89^ and TSS-seq data, as previously described^19^.

### RNA-seq library processing

The random hexamer molecular barcode on the 5′ end of each read was removed and processed using a custom Python script. Cutadapt^90^ was then used to remove adapter sequences and low-quality base calls from the 3′ end of the read, as well as poly-A sequences from the 5′ end of the read. Reads were aligned to the combined *S. cerevisiae* and *S. pombe* genome using TopHat2^91^ without a reference transcriptome, and uniquely mapping alignments were selected using SAMtools^92^. Alignments mapping to the same location as another alignment with the same molecular barcode were identified as PCR duplicates and removed using a custom Python script. Coverage corresponding to the 3′-most base of the RNA fragment sequenced as well as the entire interval for each alignment was extracted using BEDTools^93^ genomecov and normalized to the total number of alignments uniquely mapping to the *S. pombe* genome. The quality of raw, cleaned, non-aligning, and uniquely aligning non-duplicate reads was assessed using FastQC^94^.

### *Ab initio* transcript annotation from RNA-seq data

StringTie^95^ was used to perform *ab initio* transcript annotation for each RNA-seq sample, with minimum transcript length of 50 bp, minimum transcript coverage of 2.5, and minimum separation between transcripts of 12 nt. Transcript annotations for all samples (two replicates each of DMSO- and IAA-treated wild type, Spn1-noAID, and Spn1-AID samples) were merged, and those annotations not overlapping known transcripts were combined with the annotations of known transcripts to generate a unified transcript annotation. Transcripts from this list were classified as antisense if their 5′ end overlapped a known transcript on the opposite strand.

### RNA-seq differential expression analysis

For differential expression analyses of verified coding genes, RNA-seq alignments overlapping the transcript annotation of these genes were counted using BEDTools^93^. To compare Spn1-AID treated with IAA versus DMSO, the transcript counts from these two conditions were used to perform differential expression analysis using DESeq2^96^ at a false discovery rate of 0.1. All differential expression analyses were normalized to the spike-in control by using size factors obtained from *S. pombe* transcript counts for each sample. To obtain changes in verified coding transcript abundance due to Spn1 depletion while accounting for possible effects from expressing the TIR1 ubiquitin ligase, AID-tagging Spn1, or treating yeast cells with IAA, a separate multi-factor differential expression analysis was performed (Supplementary Fig. 2a). Transcript counts from all RNA-seq samples (two replicates each of DMSO- and IAA-treated wild type, Spn1-noAID, and Spn1-AID strains) were used to perform a multi-factor differential expression analysis using DESeq2^96^ at a false discovery rate of 0.1. The design formula for the multi-factor analysis was a generalized linear model with no interaction terms, in which the variables represented the presence of TIR1, the presence of the AID tag on Spn1, the presence of IAA in the media, and the depletion of Spn1. To obtain changes for all annotated transcripts, including antisense transcripts (Supplementary Fig. 2e), a multi-factor differential expression analysis of the same design was performed using counts measured over the unified transcript annotation described above.

### Gene set enrichment analysis

The set of genes with altered expression during the environmental stress response (ESR)^43^ was separated into ESR-upregulated and ESR-downregulated genes based on the median fold change of expression across stress conditions. Gene set enrichment analysis^97^ was performed for these gene sets as well as for gene ontology sets curated by the Saccharomyces Genome Database^98^, with genes ranked by their RNA-seq fold-change in the Spn1-AID strain treated with IAA versus DMSO.

### Intron retention analysis

Intron retention was defined as the proportion of unspliced RNA for an intron and was estimated from counts of RNA-seq alignments. Spliced and unspliced alignments were counted for 252 non-mitochondrial introns not overlapping 5′ UTRs^87^. Alignments overlapping an intron on the same strand were counted as spliced if the alignment indicated matches of at least 4 nt with both the 5′ and 3′ exon sequences, separated by a skipped region over the coordinates of the intron. Alignments overlapping an intron on the same strand were counted as unspliced if the alignment indicated a continuous match spanning at least 4 nt on both sides of the 5′ or 3′ splice junction, or if the alignment was contained entirely within the intron boundaries. Alignments overlapping annotated features contained within intron boundaries were excluded to avoid conflating differential expression of these features with changes in intron retention. Spliced and unspliced alignment counts were pooled between replicates. To address high variance in the raw intron retention proportion 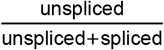 associated with low coverage introns, intron retention was estimated using empirical Bayes shrinkage towards a beta-binomial prior. The prior distribution was fit by maximum likelihood estimation using the intron alignment counts for the 217/252 introns with at least one spliced alignment in any of the conditions tested. To determine the significance of changes in intron retention upon Spn1 depletion, Monte Carlo sampling from the posterior intron retention distributions was used to determine the probability that intron retention in the Spn1-AID strain treated with IAA was more extreme than intron retention in all control conditions (wild type DMSO, wild type IAA, Spn1-noAID DMSO, Spn1-noAID IAA, and Spn1-AID DMSO).

### ChIP-seq library processing

Reads were demultiplexed using fastq-multx^99^, allowing one mismatch to the index sequence and A-tail. Cutadapt^90^ was then used to remove index sequences and low-quality base calls from the 3′ end of the read. Reads were aligned to the combined *S. cerevisiae* and *S. pombe* genome using Bowtie 2^100^, and alignments with a mapping quality of at least 5 were selected using SAMtools^92^. The median fragment size estimated by MACS2^101^ cross-correlation over all samples of a factor was used to generate coverage of fragments and fragment midpoints by extending alignments to the median fragment size or by shifting the 5′ end of alignments by half the median fragment size, respectively. The quality of raw, cleaned, non-aligning, and uniquely aligning reads was assessed using FastQC^94^.

### ChIP-seq normalization

For ChIP-seq coverage from IP samples, spike-in normalization was accomplished by scaling coverage proportionally to the normalization factor 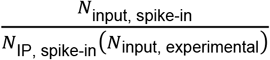, where *N*_IP, spikein_ is the number of *S. pombe* alignments in the IP sample, *N*_input, spike-in_ is the number of *S. pombe* alignments in the corresponding input sample, and *N*_input, experimental_ is the number of *S. cerevisiae* alignments in the input sample. Coverage from input samples was normalized to *N*_input, experimental_. Relative estimates of the total abundance of each ChIP target on chromatin were also obtained by multiplying the normalization factor with the number of *S. cerevisiae* alignments in an IP sample. Coverage of the relative ratio of IP over input was obtained by first smoothing normalized IP and input fragment midpoint coverage using a Gaussian kernel with 20 bp bandwidth, and then taking the ratio. Coverage of the relative ratio of one factor to another (e.g. H3K36me3 over H3) was obtained as follows: For each factor, coverage of IP over input in each sample was standardized using the genome-wide mean and standard deviation over all samples, weighted such that the non-depleted and Spn1-depleted conditions had equal contribution. In the cases where ChIP for both factors was performed using the same chromatin preparation, the standardized coverage of the normalizing factor was subtracted from the matched coverage of the factor to be normalized. In the cases where ChIP for each factor was performed using different chromatin preparations (Spt6 and Set2 normalized to Rpb1), standardized coverages were averaged across replicates before subtracting coverage of the normalizing factor from the coverage of the factor to be normalized.

### ChIP-seq differential occupancy analysis

For differential occupancy analyses of single factors over verified coding genes, IP and input fragment midpoints overlapping the transcript annotation of these genes were counted using BEDTools^93^. These counts were used to perform a differential occupancy analysis using DESeq2^96^, at a false discovery rate of 0.1. The design formula used was a generalized linear model with variables for sample type (IP or input), condition (non-depleted or Spn1-depleted), and the interaction of sample type with condition. Fold changes were extracted from the coefficients of the interaction of sample type with condition, and represent the change in IP signal observed upon Spn1 depletion, corrected for changes in input signal. When normalizing to the spike-in control, size factors obtained from *S. pombe* counts over peaks called with MACS2^101^ were used for each sample. All single-factor differential occupancy results shown are normalized to spike-in, except for H3 differential occupancy results, where the additional variability introduced by normalization to spike-in would have complicated identification of the set of genes with significantly reduced H3. For differential occupancy analysis of H3 over the first 500 bp of verified coding genes (Supplementary Fig. 4b), the same approach was taken except using counts over the relevant regions. For differential occupancy analyses considering one factor normalized to another (e.g. changes in Rpb1-normalized Spt6), IP and input counts for both factors were used to perform a differential binding analysis using DESeq2^96^, at a false discovery rate of 0.1. The design formula used was a generalized linear model with all crossings of variables for sample type (IP or input), ChIP target, and condition (non-depleted or Spn1-depleted). Fold changes were extracted from the coefficients of the interaction of sample type, ChIP target, and condition. These values represent changes in input-normalized IP signal of one factor upon Spn1 depletion, corrected for changes in input-normalized IP signal of the other factor. All factor-normalized differential occupancy analysis results shown are normalized to spike-in by using size factors obtained from counts over *S. pombe* transcripts.

## Reporting summary

Further information on research design is available in the Nature Research Reporting Summary linked to this article.

## Data availability

The RNA-seq and ChIP-seq data sets are available in the GEO repository, accession number GSE138281. They can be accessed at https://www.ncbi.nlm.nih.gov/geo/query/acc.cgi?acc=GSE153567. All other relevant data supporting the key findings of this study are available within the article and its Supplementary Information files or from the corresponding authors upon reasonable request. A reporting summary for this article is available as a Supplementary Information file.

## Code availability

An archive including all code used is available at Zenodo (doi.org/10.5281/zenodo.3901642).

## Acknowledgments

We thank Olga Viktorovskaya for suggesting and helping with the co-immunoprecipitation experiments. We thank Raja Gopalakrishnan, Olga Viktorovskaya, and James Warner for helpful comments on the manuscript. N.I.R. was supported by F31 GM112370. This work was supported by NIH grant R01GM120038 to F.W.

## Author contributions

N.I.R. and F.W. designed the experiments. N.I.R. performed all the experiments. J.C., D.J., and B.A. performed computational analyses with support from P.J.P. for D.J., and B.A. N.I.R., J.C., and F.W. wrote the paper.

## Additional information

Correspondence and requests for materials should be addressed to F.W.

---

**Supplementary Table 1.**
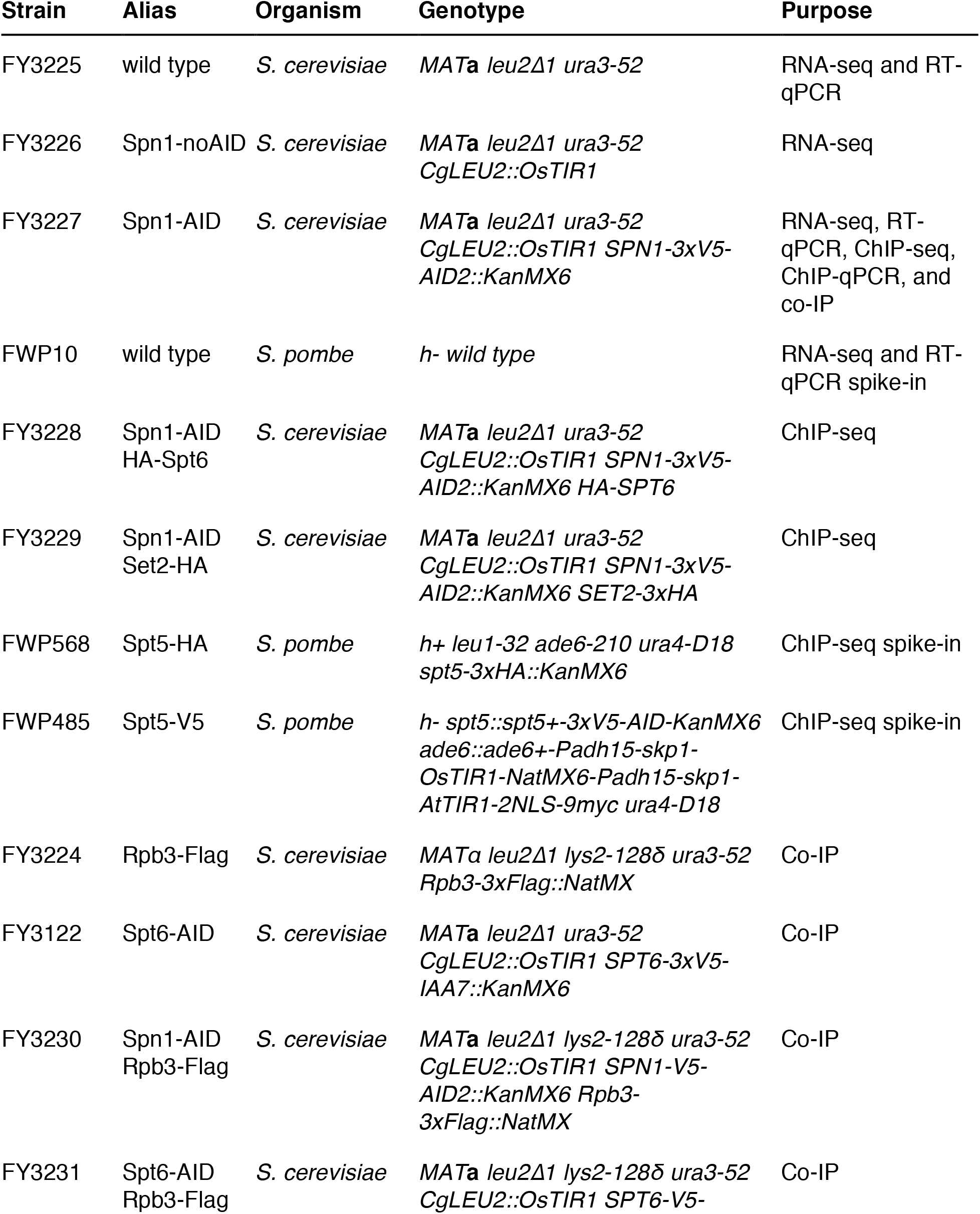
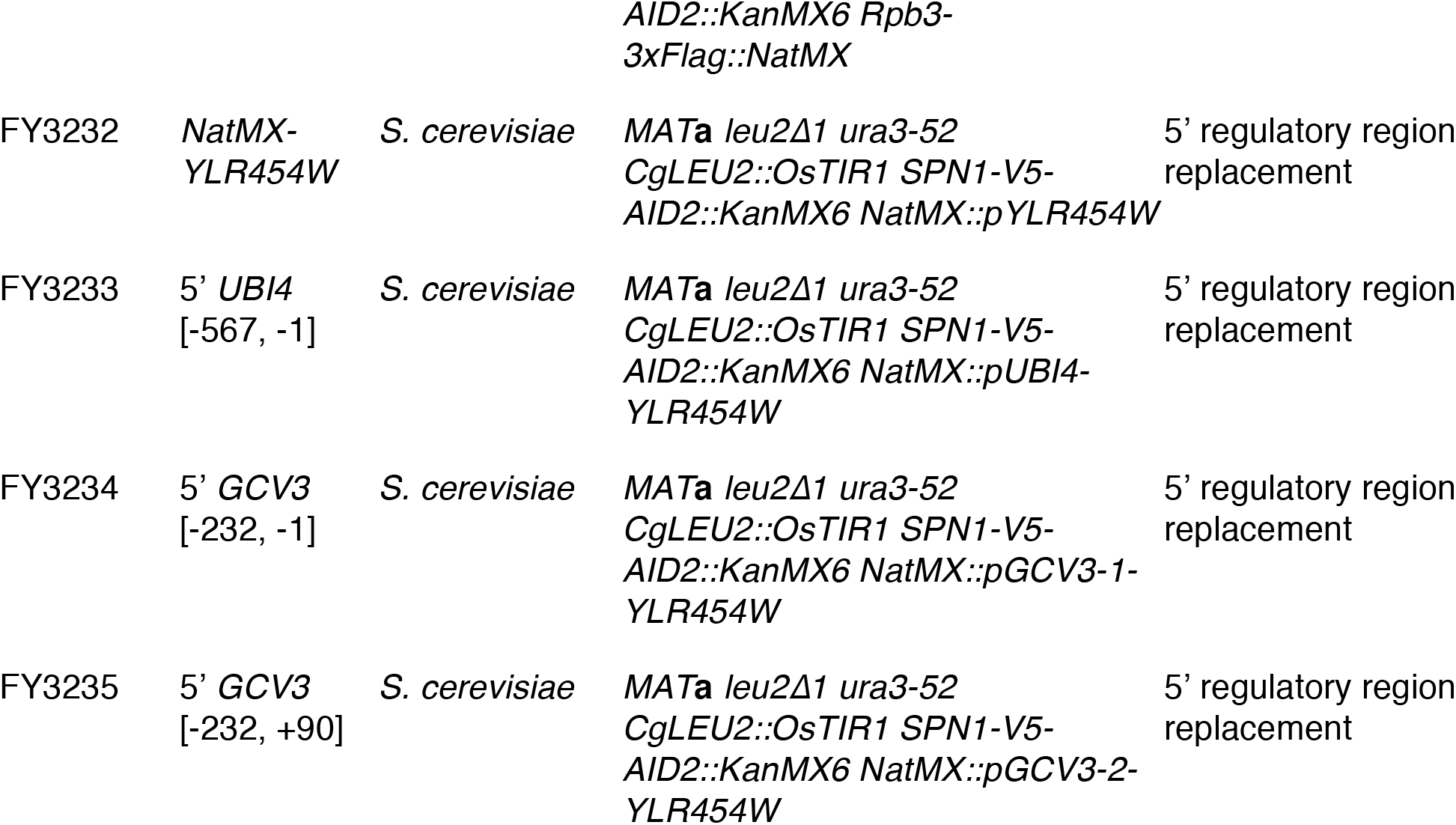
Yeast strains used in this study Strain Alias Organism Genotype.

**Supplementary Table 2.**
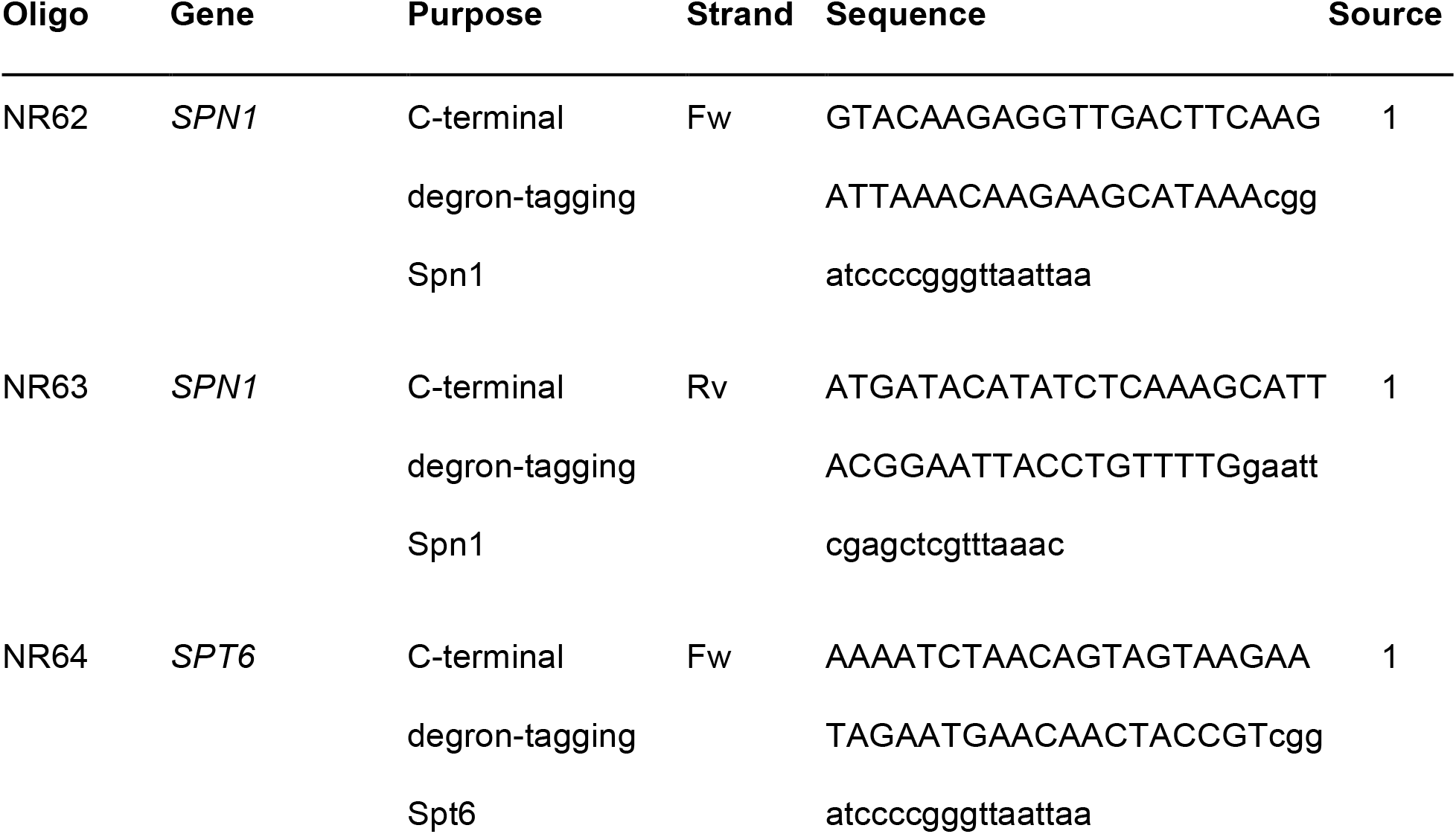
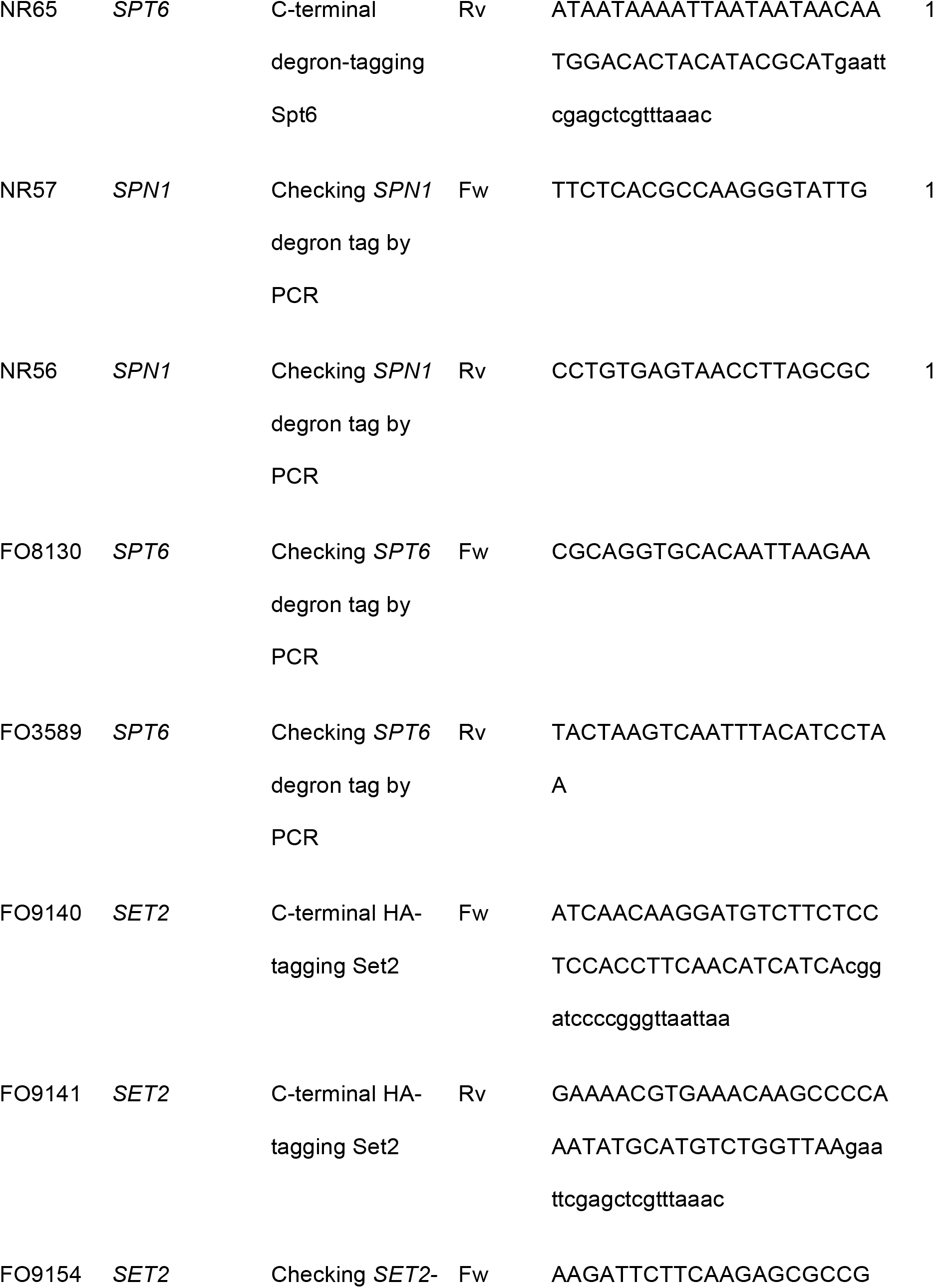
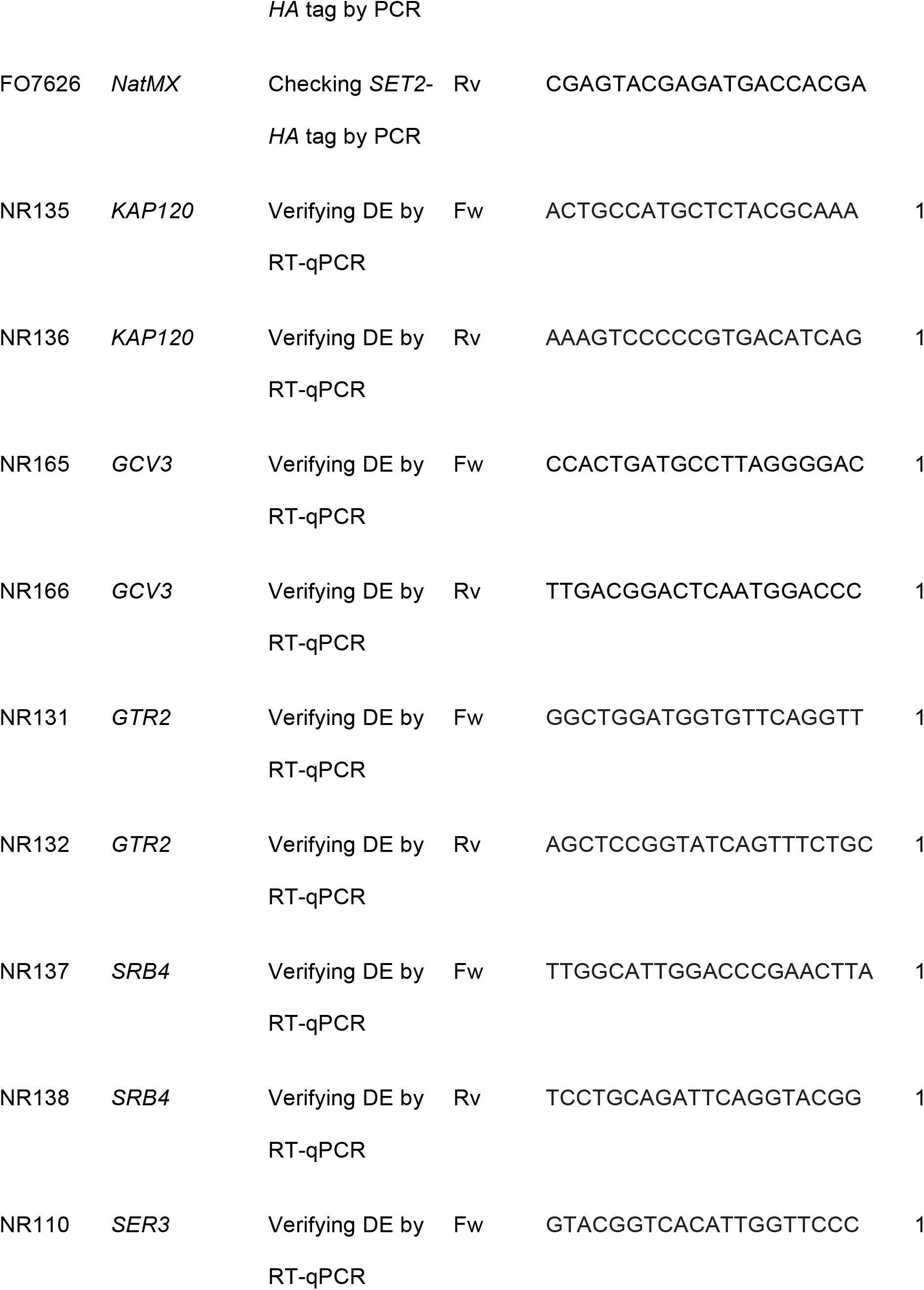
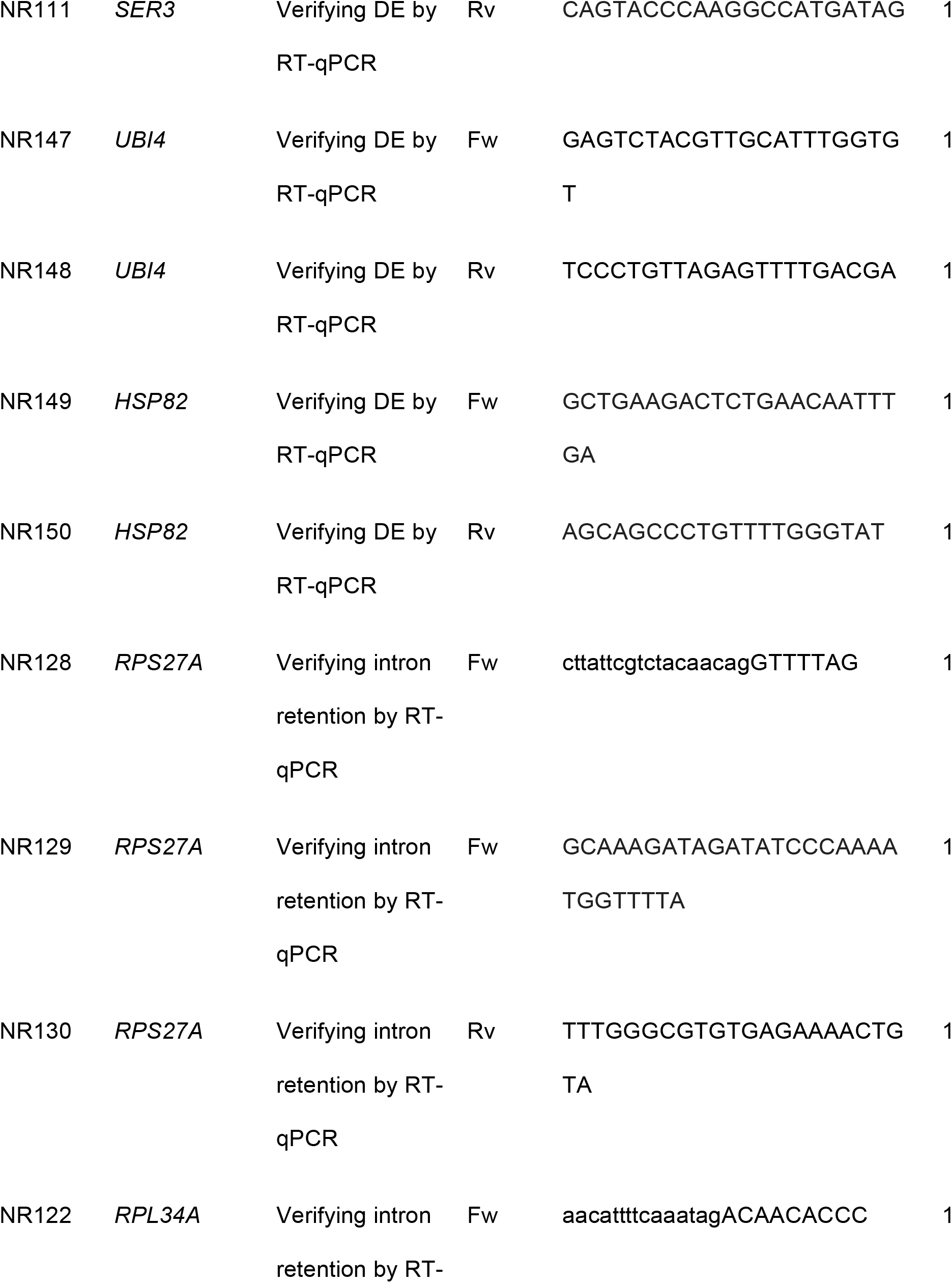
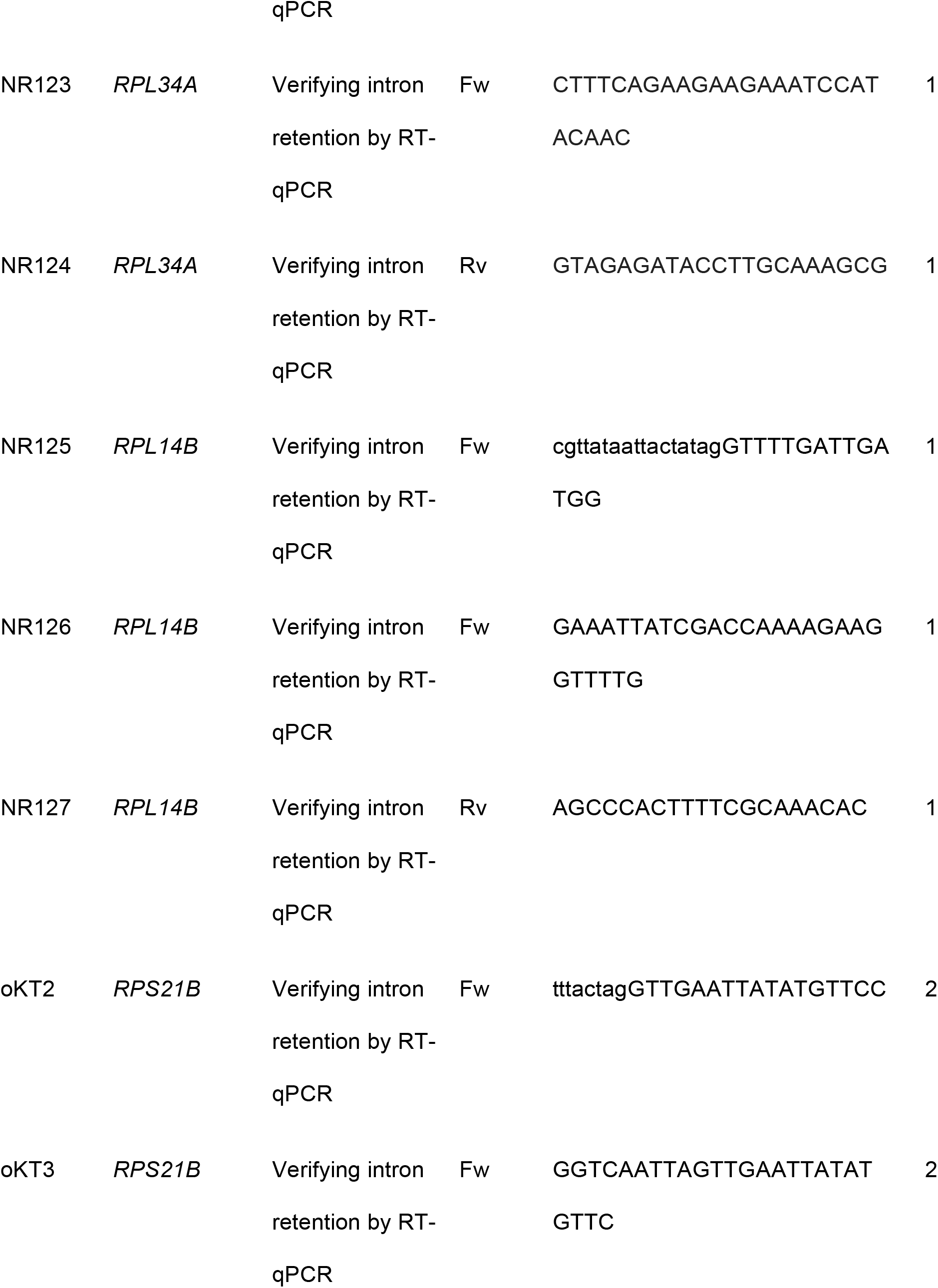
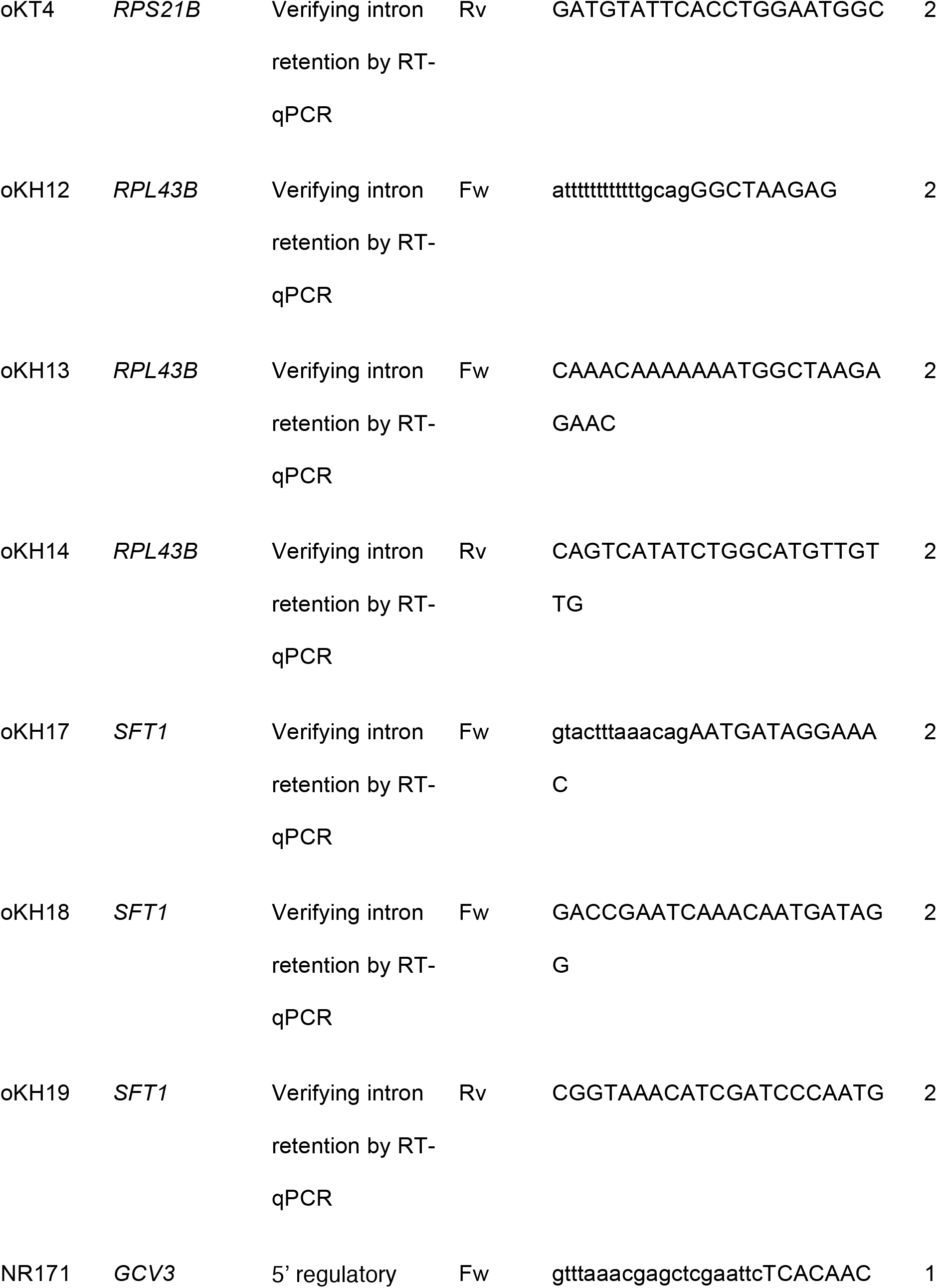
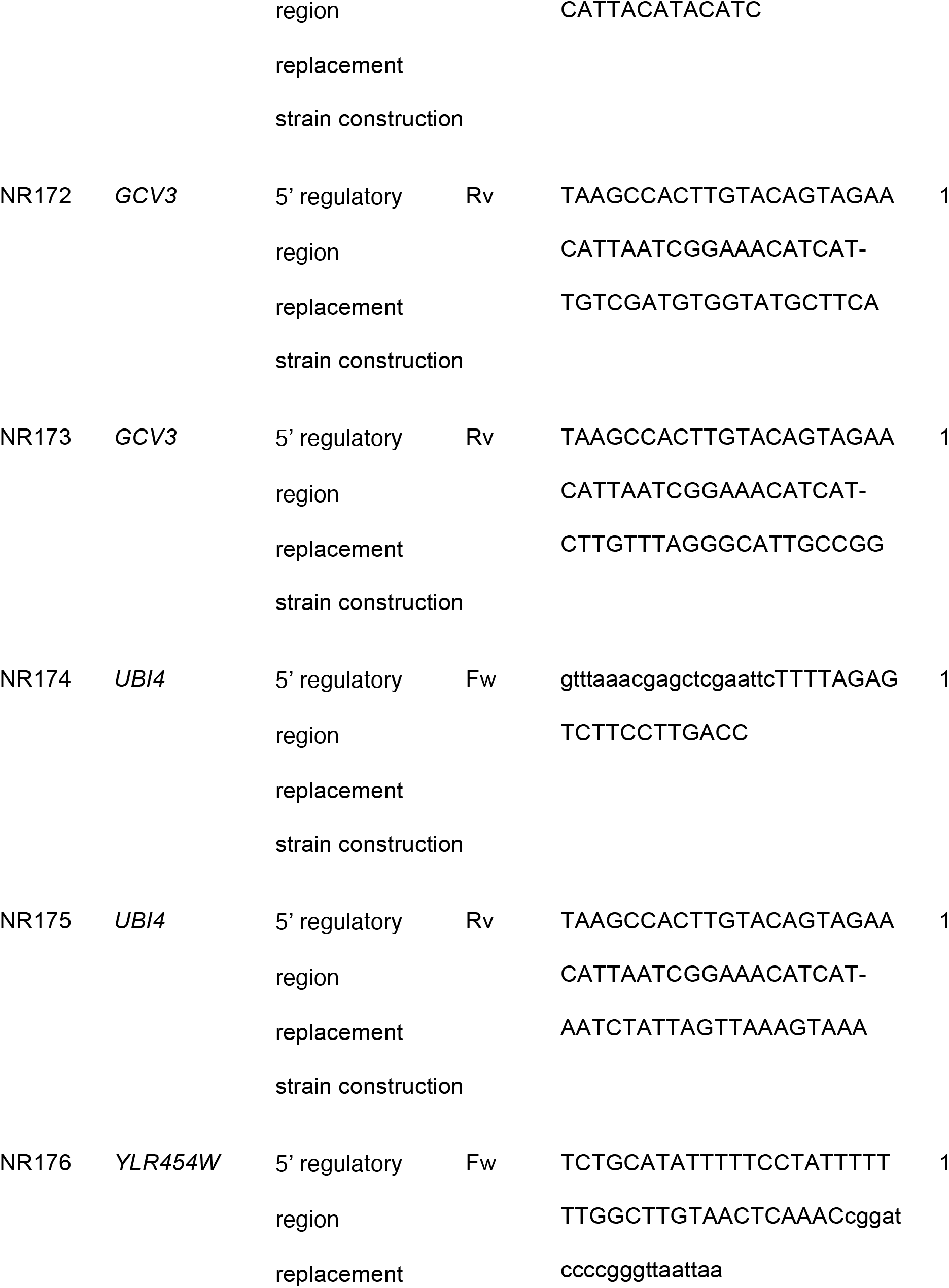
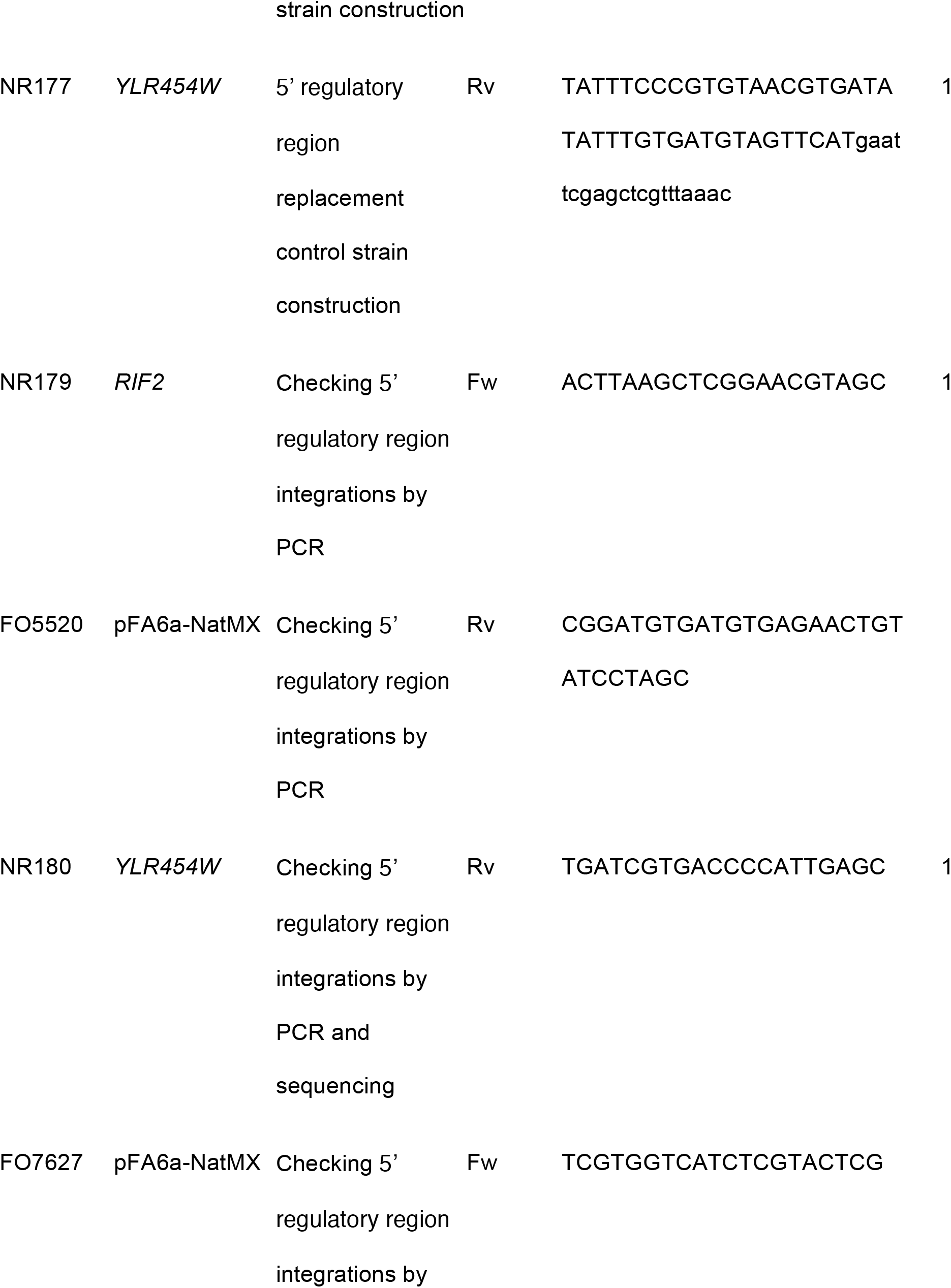
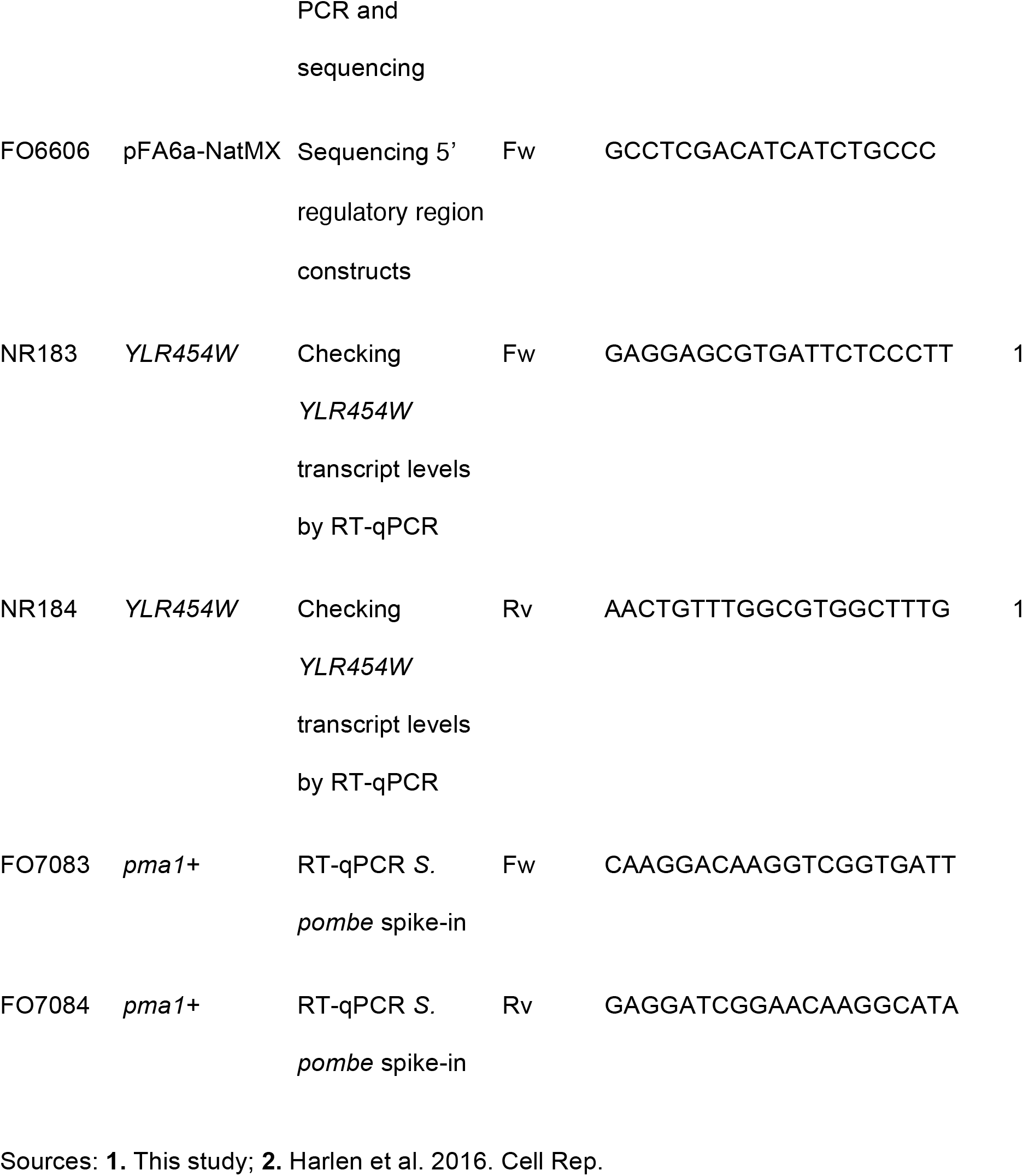
Primers used in this study.

**Supplementary Figure 1.**
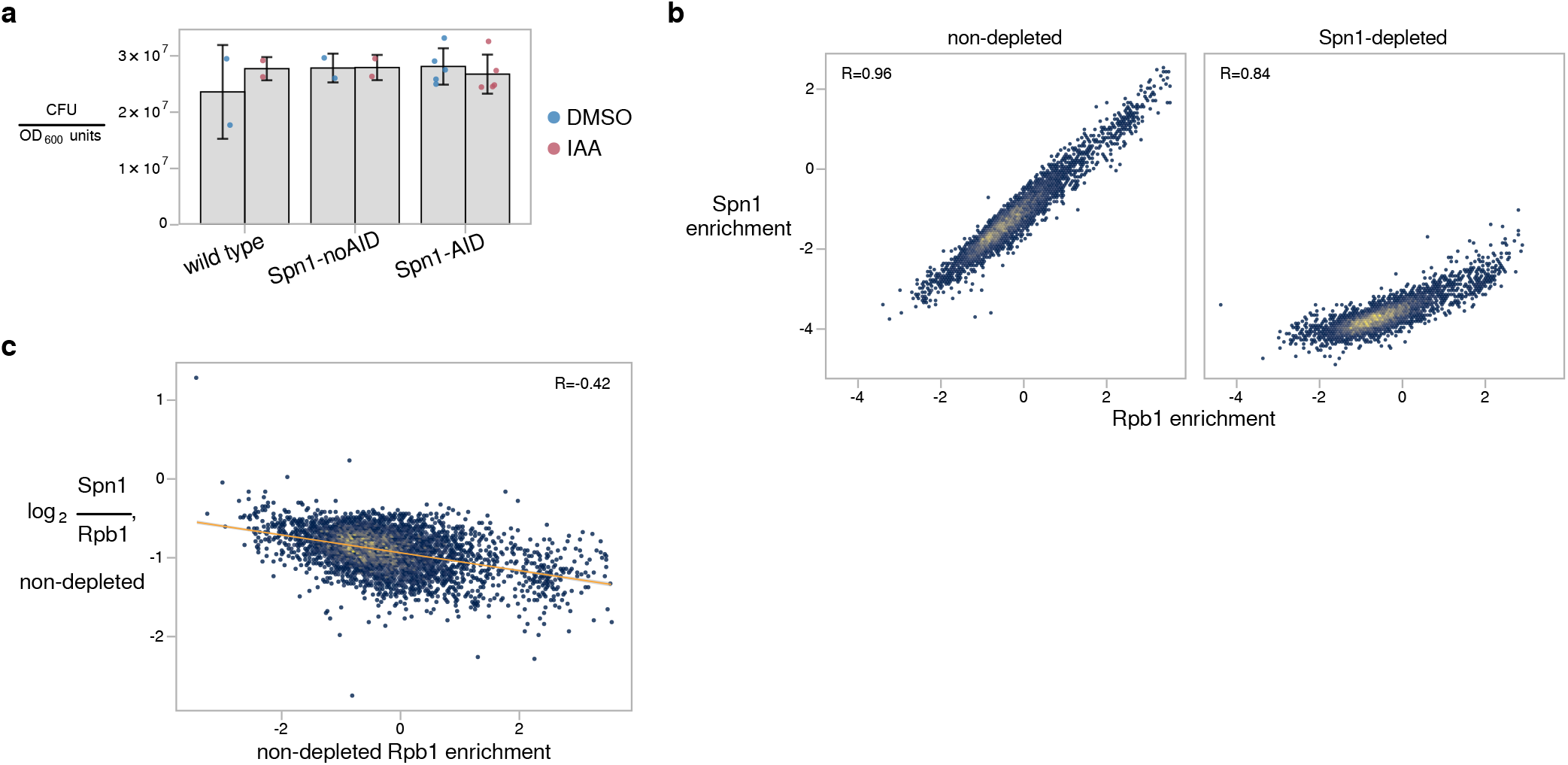
Effects of Spn1 depletion. **a** Viability of a wild-type strain, a strain expressing TIR1 but lacking the degron tag on Spn1 (Spn1-noAID), and the Spn1 degron strain (Spn1-AID) after 90 minutes of treatment with DMSO or 25 μM IAA, quantified as colony-forming units per OD_600_ unit of culture plated. Error bars indicate the mean ± standard deviation of the replicates shown. **b** Scatterplots of Spn1 enrichment versus Rpb1 enrichment for 5,091 verified coding genes in non-depleted and Spn1-depleted conditions. Pearson correlation coefficients are indicated. Enrichment values are the relative log_2_ enrichment of IP over input. **c** Scatterplot showing Rpb1-normalized Spn1 enrichment versus Rpb1 enrichment for the same genes as in (b) in the non-depleted condition. Rpb1 enrichment values are the relative log_2_ enrichment of IP over input. The Pearson correlation coefficient is indicated, and a linear regression is overlaid.

**Supplementary Figure 2.**
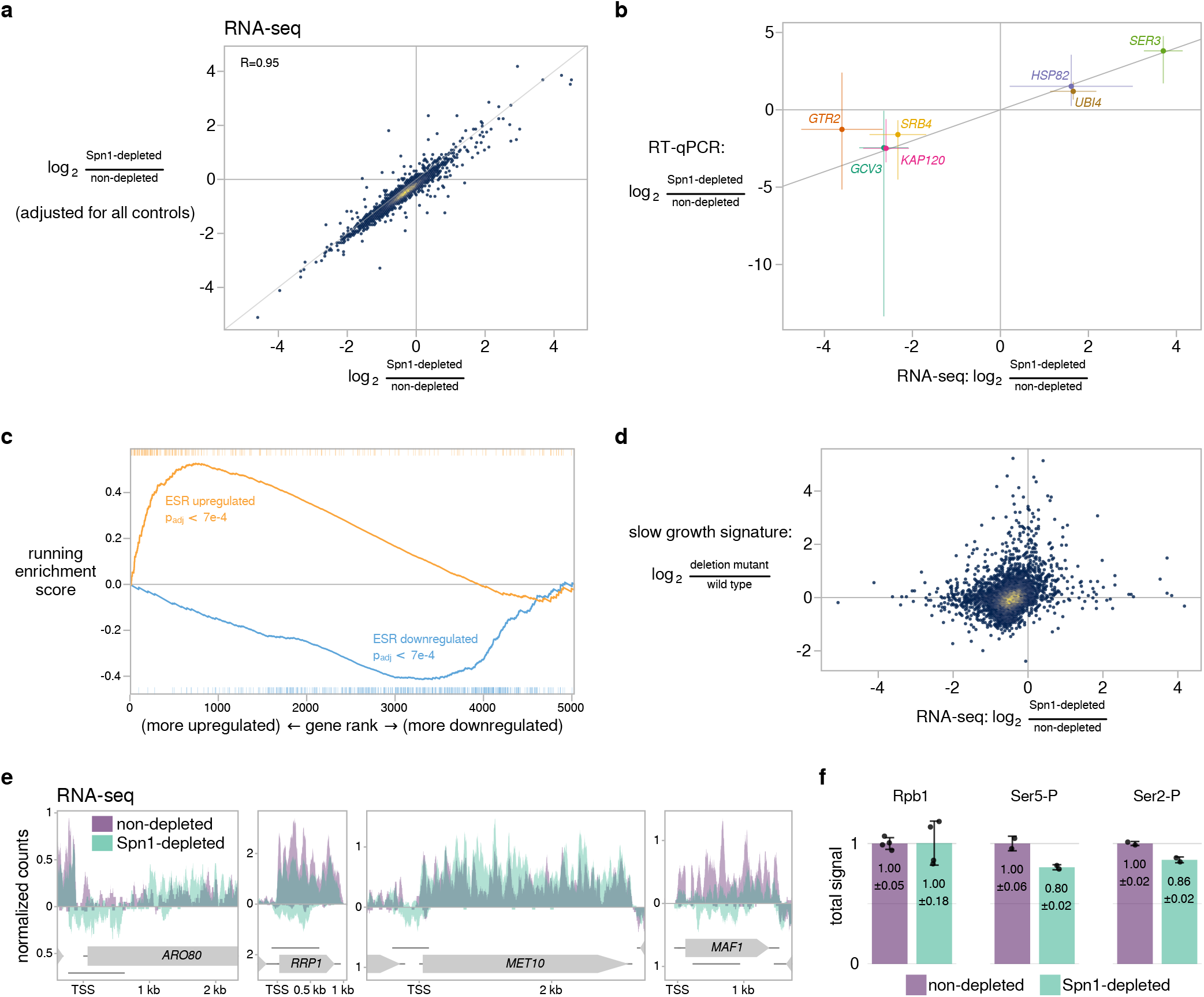
RNA-seq and RNAPII ChIP-seq. **a** Scatterplot comparing the results of two RNA-seq differential expression analyses for 5,091 verified coding genes. X-axis values show the fold-change estimated from comparison of the Spn1 degron strain treated with IAA (Spn1-depleted) versus DMSO (non-depleted), the same comparison shown in Fig. 2a. Y-axis values show the fold-change attributable to the effects of Spn1 depletion, as extracted from a multi-factor analysis incorporating RNA-seq data from additional control strains to be able to separate effects due to Spn1 depletion from effects due to presence of the degron tag on Spn1, expression of degron system component TIR1, or treatment of yeast cells with IAA. The line y=x is overlaid for comparison. Results of the simple comparison between Spn1-depleted and non-depleted conditions are used throughout this study for consistency with ChIP-seq analyses for which the multi-factor analysis was not performed. **b** Scatterplot comparing changes in transcript abundance upon Spn1 depletion as measured by RT-qPCR and RNA-seq. The line y=x is drawn for comparison, and error bars indicate 95% confidence intervals. **c** Running GSEA enrichment score for genes up- and downregulated during the yeast environmental stress response (ESR)43 along genes ranked by change in transcript abundance upon Spn1 depletion as measured by RNA-seq. The ranks of ESR up- and downregulated genes are marked along the top and bottom of the plot, respectively, and Benjamini-Hochberg adjusted enrichment p-values are indicated. **d** Scatterplot comparing changes in transcript abundance upon Spn1 depletion as measured by RNA-seq versus changes in transcript abundance expected from a slow growth signature common to many environmental and genetic perturbations as previously reported^44^. **e** Spike-in normalized RNA-seq coverage over four genes with antisense transcripts that are differentially upregulated upon Spn1 depletion. Sense and antisense strand signal are plotted above and below the x-axis, respectively. Unlabeled horizontal lines indicate the boundaries of transcripts called by StringTie. **f** Barplots showing total levels of Rpb1, Rpb1-Ser5-P, and Rpb1-Ser2-P on chromatin in non-depleted and Spn1-depleted conditions, estimated from ChIP-seq spike-in normalization factors (see Methods). The values and error bars for each condition indicate the mean ± standard deviation of the replicates shown. Note that we observed batch-to-batch variability in Rpb1 levels after Spn1 depletion, with Rpb1 levels decreasing for the two replicates generated from the same chromatin preparations used to ChIP Rpb1-Ser5-P and Rpb1-Ser2-P.

**Supplementary Figure 3.**
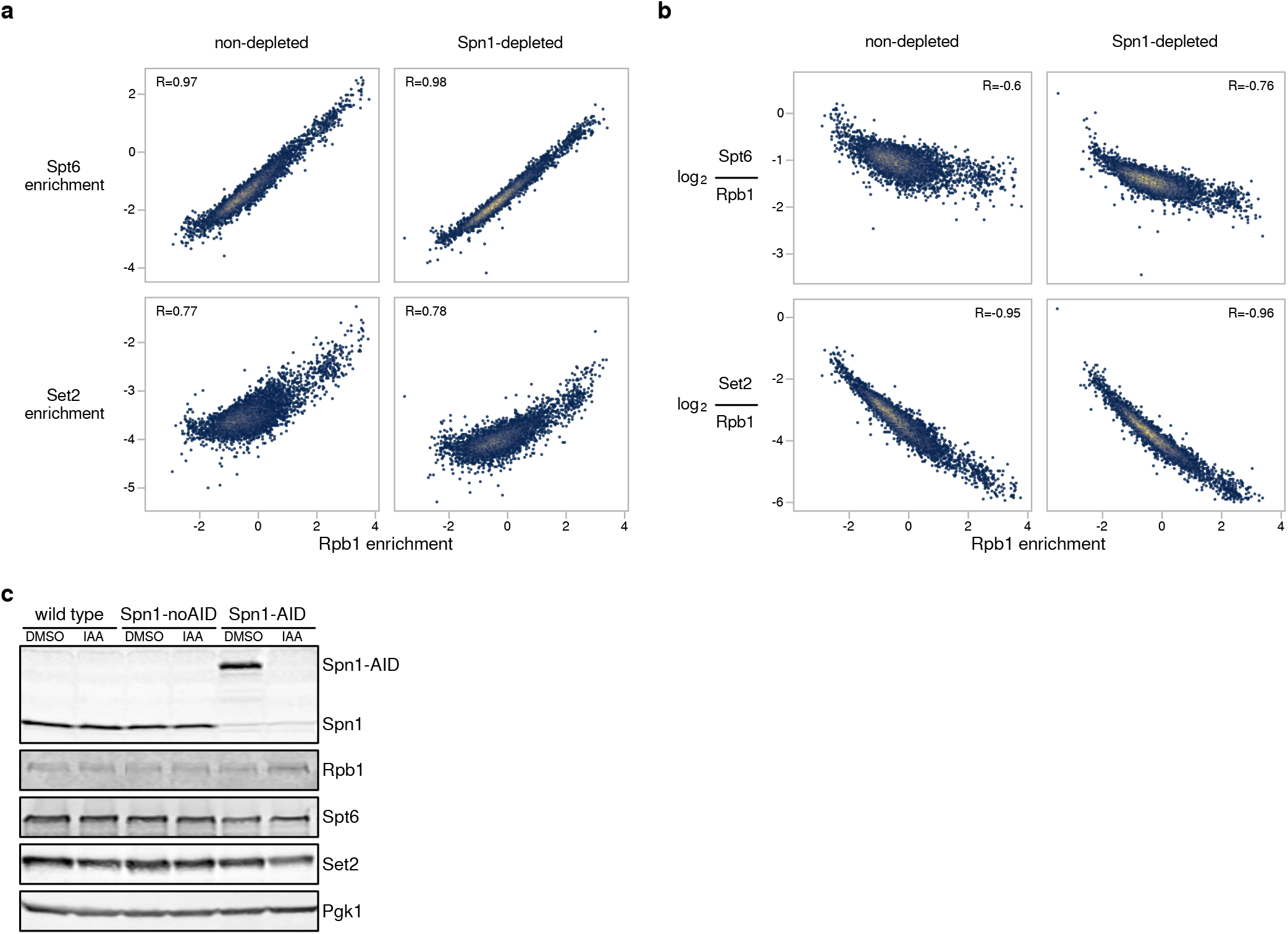
Spt6 and Set2 ChIP-seq. **a** Scatterplots showing Spt6 or Set2 enrichment versus Rpb1 enrichment for 5,091 verified coding genes in non-depleted and Spn1-depleted conditions. Enrichment values are the relative log_2_ enrichment of IP over input. Pearson correlation coefficients are indicated. **b** Scatterplots showing Rpb1-normalized enrichment of Spt6 or Set2 versus Rpb1 enrichment for the same genes as in (a) in non-depleted and Spn1-depleted conditions. Pearson correlation coefficients are indicated. **c** Representative western blots showing the levels of Spn1, Rpb1, Spt6, and Set2 in the wild type, Spn1-noAID, and Spn1-AID strains treated with DMSO or 25 μM IAA. The faint band co-migrating with native Spn1 in the Spn1-AID lanes of the Spn1 blot is non-specific, as it was also present in a *spn1*Δ strain (data not shown). Pgk1 was used as a loading control.

**Supplementary Figure 4.**
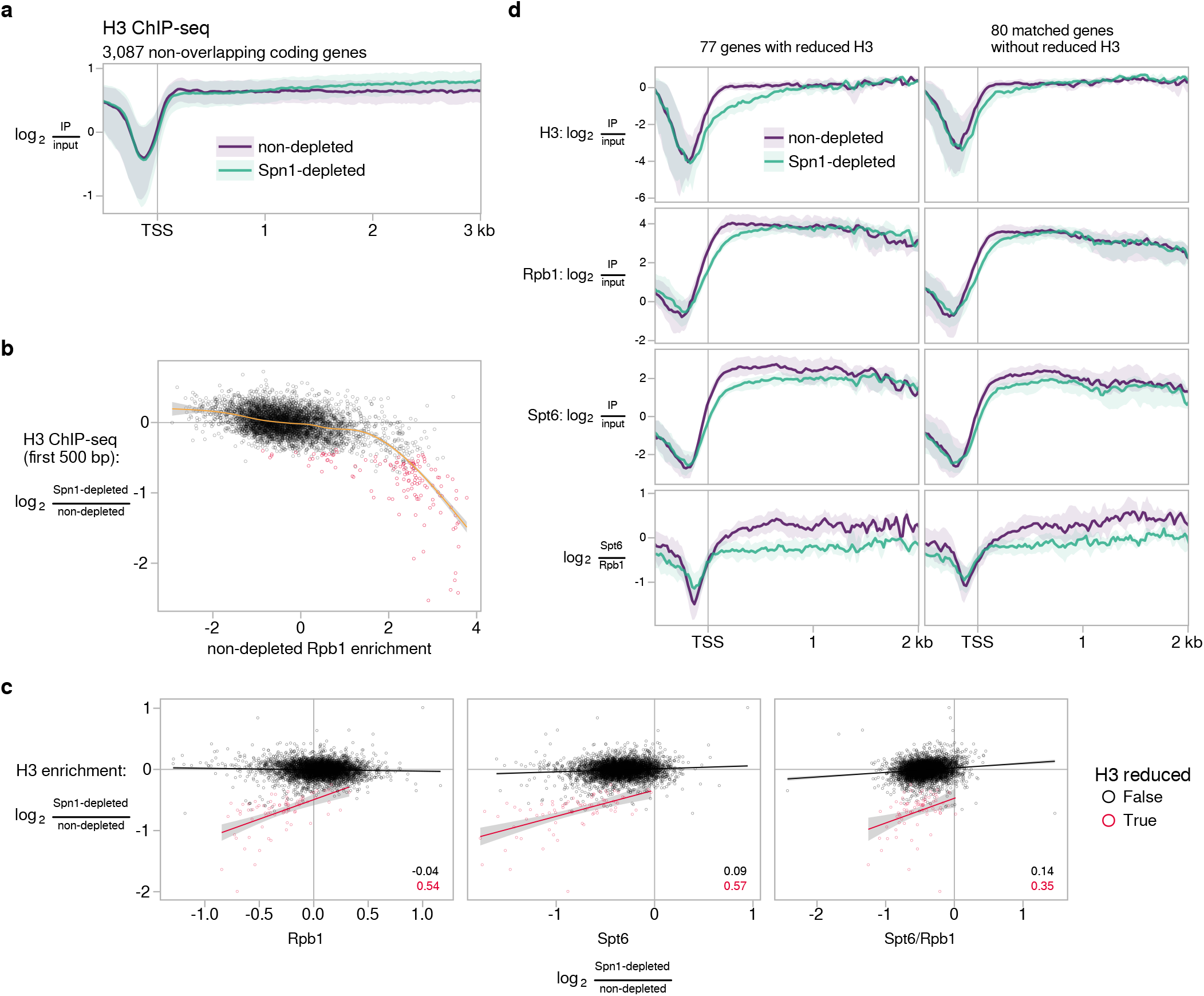
Histone H3 ChIP-seq. **a** Average H3 ChIP enrichment for 3,087 non-overlapping, verified coding genes aligned by TSS in non-depleted and Spn1-depleted conditions. The solid line and shading are the median and interquartile range of the mean enrichment over four replicates. **b** Scatterplot showing change in H3 enrichment upon Spn1 depletion over the first 500 bp downstream of the TSS versus non-depleted Rpb1 enrichment over the entire length of 4,924 verified coding genes at least 500 bp long. Rpb1 enrichment values are the relative log_2_ enrichment of IP over input. Genes with significant decreases in H3 enrichment over their first 500 bp are colored red, and a cubic regression spline is overlaid. **c** Scatterplots comparing Spn1-dependent changes in H3 enrichment over the entire gene to changes in Rpb1 enrichment, Spt6 enrichment, and Rpb1-normalized Spt6 enrichment. Values are shown for 5,091 verified coding genes, with genes for which H3 enrichment is significantly reduced upon Spn1 depletion colored red. A linear regression for each group is overlaid, with Pearson correlation coefficients shown in the bottom right. **d** Average H3, Rpb1, Spt6, and Rpb1-normalized Spt6 ChIP enrichment in non-depleted and Spn1-depleted conditions for the 77 genes with significantly reduced H3 enrichment upon Spn1 depletion and for 80 genes without significantly reduced H3 enrichment matched to the first group by expression level and transcript length. The solid line and shading are the median and interquartile range over the genes in each group.

**Supplementary Figure 5.**
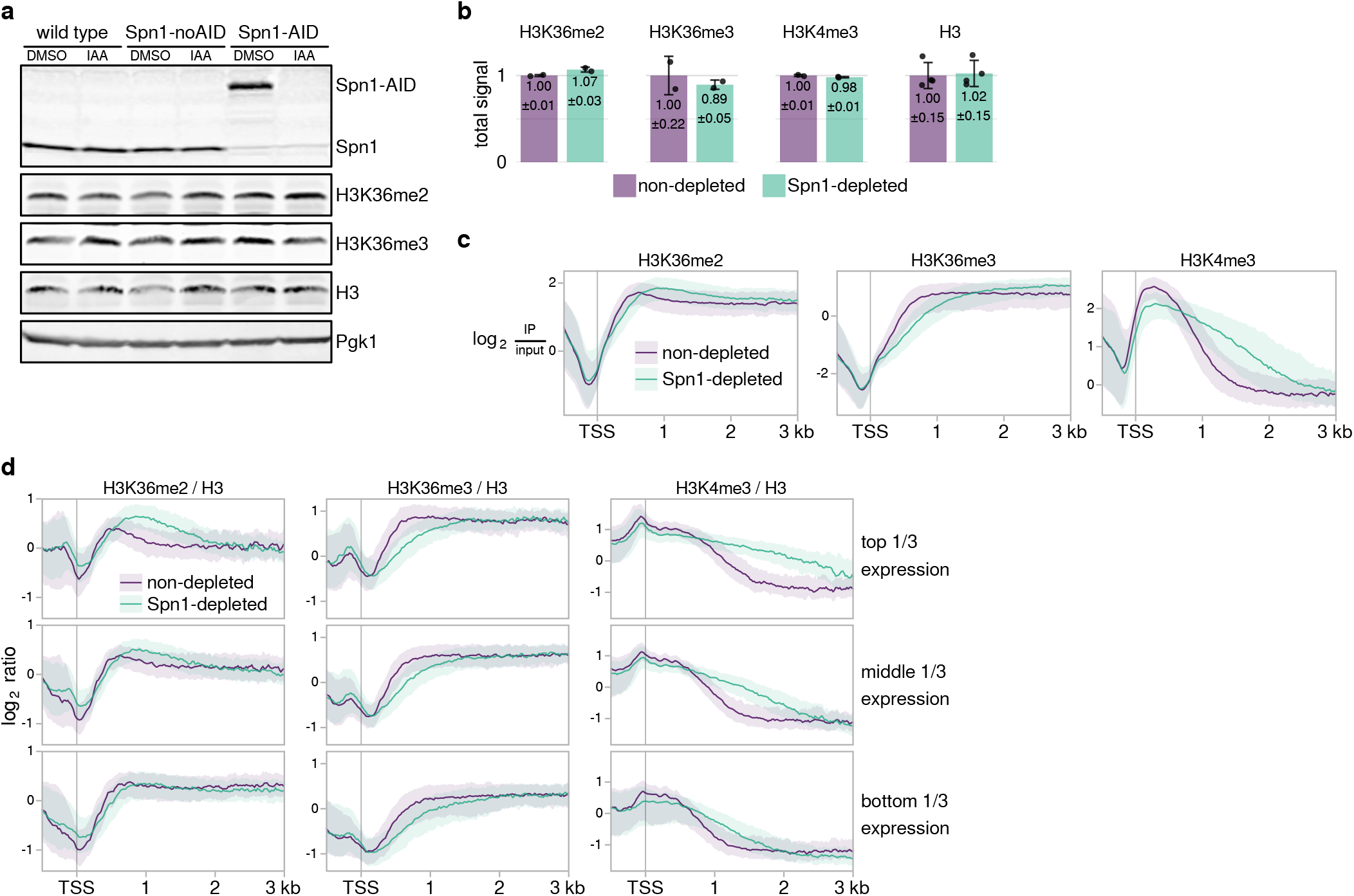
Histone H3 modifications ChIP-seq. **a** Representative western blots showing the levels of H3K36me2, H3K36me3, and H3 in the wild type, Spn1-noAID, and Spn1-AID strains treated with DMSO or 25 μM IAA. The same Spn1 and Pgk1 panels used in Supplementary Fig. 3c are shown here for reference, as they were generated as part of the same experiment. **b** Barplots showing total levels of H3K36me2, H3K36me3, H3K4me3, and H3 on chromatin in non-depleted and Spn1-depleted conditions, estimated from ChIP-seq spike-in normalization factors (see Methods). The values and error bars for each condition indicate the mean ± standard deviation of the replicates shown. **c** Average H3K36me2, H3K36me3, and H3K4me3 ChIP enrichment in non-depleted and Spn1-depleted conditions for 3,087 non-overlapping, verified coding genes aligned by TSS. The solid line and shading are the median and interquartile range of the mean enrichment over two replicates. **d** Average H3-normalized H3K36me2, H3K36me3, and H3K4me3 ChIP enrichment in non-depleted and Spn1-depleted conditions for 3,087 non-overlapping, verified coding genes separated into tertiles of expression by their non-depleted Rpb1 ChIP enrichment. The solid line and shading are the median and interquartile range of the mean ratio over two replicates.

**Supplementary Figure 6.**
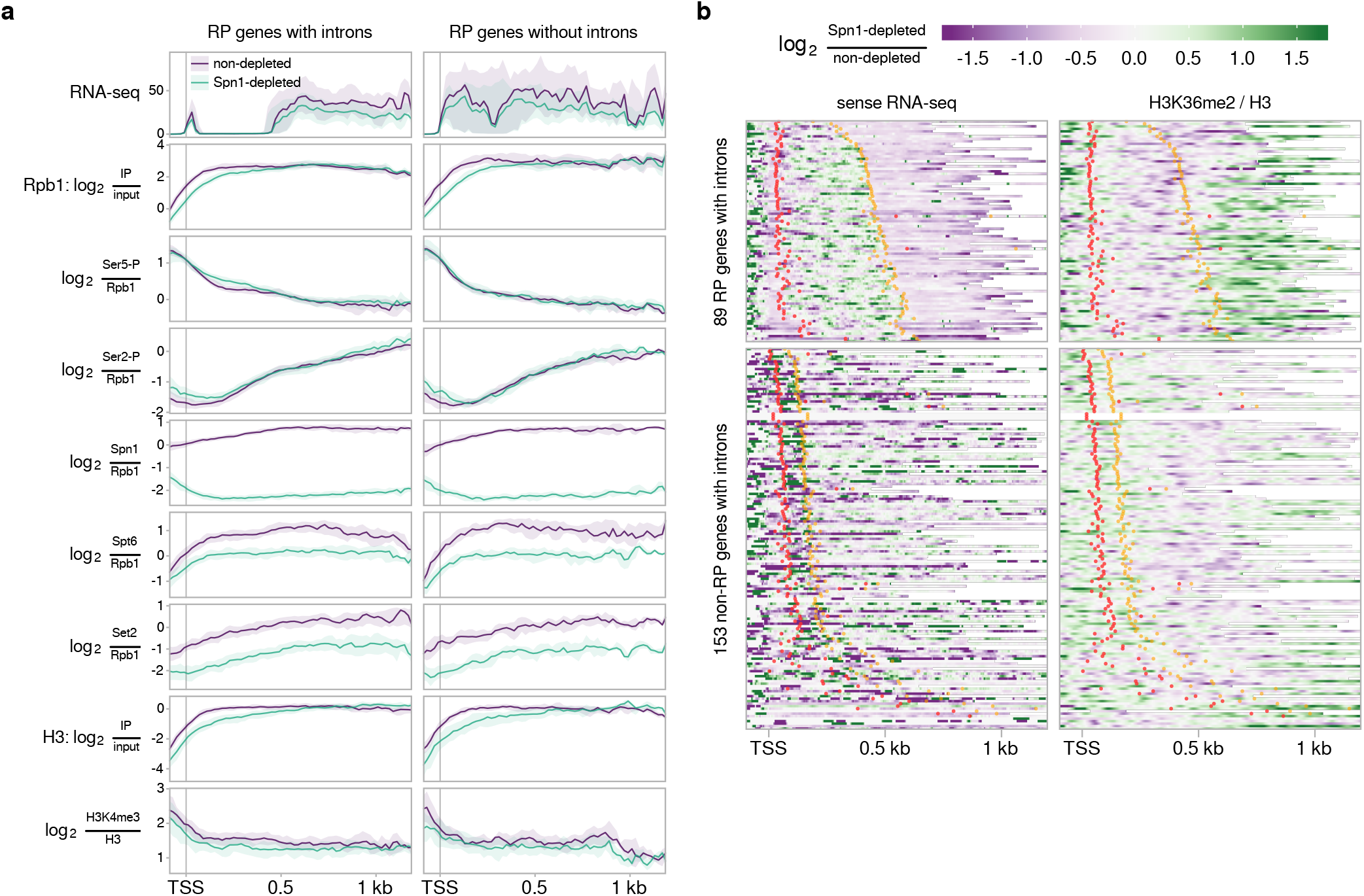
Effects of Spn1 depletion at intron-containing ribosomal protein genes. **a** Genomic data visualization for ribosomal protein genes with and without introns. The solid line and shading for each assay are the median and interquartile range over the genes in each group. The same sense strand RNA-seq coverage panel used in Fig. 7c is shown here for reference. **b** Heatmaps of change upon Spn1 depletion for sense strand RNA-seq signal and H3-normalized H3K36me2 ChIP enrichment over 89 ribosomal protein (RP) genes with introns and 153 non-RP genes with introns. Genes are aligned by TSS and ordered by distance from the TSS to the midpoint of the first intron. The 5*′* and 3*′* splice sites of each intron are marked with red and orange dots, respectively.

